# The Ndc80-Cdt1-Ska1 complex constitute a minimal processive kinetochore-microtubule coupling unit

**DOI:** 10.1101/2022.05.05.490787

**Authors:** Amit Rahi, Manas Chakraborty, Shivangi Agarwal, Kristen Vosberg, Shivani Agarwal, Annie Y. Wang, Richard J. McKenney, Dileep Varma

## Abstract

The Ndc80 kinetochore complex is essential for robust kinetochore-microtubule (k-MT) attachments during mitosis. Ndc80 has been shown to recruit the Ska1 complex to kinetochores, where Ska1 is thought to aid in k-MT coupling by Ndc80. Our previous work has shown that Cdt1, a DNA replication licensing factor, is a novel mitotic spindle-associated protein that is also recruited to kinetochores via Ndc80 and is required for stabilizing k-MT attachments. In this study, we developed auxin-induced degron (AID)-tagging to validate the previously demonstrated mitotic role of Cdt1. We demonstrate a direct interaction between Cdt1 and Ska1 that is essential for proper recruitment of Cdt1 to kinetochores and spindle microtubules. We find that Cdk1’dependent phosphorylation of Cdt1 during mitosis is critical for Ska1-binding, consequently regulating the stabilization of metaphase k-MT attachments and normal mitotic progression. Total internal reflection fluorescence microscopy (TIR-FM) experiments reveal that Cdt1 synergizes with the Ndc80 and the Ska1 complexes for microtubule-binding. Further, we show that single Cdt1 molecules form diffusive tripartite complexes with Ndc80 and Ska1 that can processively track the ends of dynamic microtubules *in vitro*. Taken together our data identifies a minimal molecular unit responsible for bidirectional processive tip tracking of kinetochores.

## Introduction

Bona-fide DNA replication is a committed step in cell proliferation that ensures complete and precise genome duplication only once per cell cycle. Cdt1 is one of the DNA replication licensing proteins that serves a central role in this process [1]. Following DNA replication, the cell undergoes mitosis to equally partition its duplicated chromosomes between the two daughter cells. This function is achieved by the concerted function of the bipolar spindle microtubules (microtubules) and kinetochores [2]. Besides licensing the origins for DNA replication, a fraction of Cdt1 has been shown to localize to the kinetochores during mitosis, dependent on the flexible loop domain of the Ndc80 complex, a key component of the core microtubule-binding site at kinetochores [3–5], where Cdt1 further augmented the stability of kinetochore-microtubule (k-MT) attachments [3, 6]. Interestingly, unlike other licensing proteins, Cdt1 lacks enzymatic activity and shares little or no resemblance to any other protein of known molecular function. Although, our earlier work demonstrating the ability of Cdt1 to bind to microtubules directly and its regulation by Aurora B Kinase-mediated phosphorylation sheds some mechanistic insights into how Cdt1 promotes stable k-MT attachments [6]; the next compelling question was howCdt1 and Ndc80 are implicated in generating stable k-MT attachments and how these activities are coordinated or synergized. This study also found that Cdt1 localized to the plus-ends of spindle microtubules when expressed in cells. *In vitro*, Cdt1 not only possessed the capacity to diffuse onmicrotubules but also was able to bind to curved microtubule protofilaments.

Interestingly, the spindle microtubule-binding Ska1 complex is another factor that localizes to kinetochores in an Ndc80-dependent manner and has been shown to assist the Ndc80 complex in the formation of load-bearing k-MT attachments in metaphase [7, 8]. On the other hand, Ndc80 was also able to strengthen the binding affinity of Ska1 to the microtubules, which is indicative that these proteins might form a complex in the presence of microtubules [9].

Based on the afore-mentioned studies, it was evident that many of the properties of Cdt1, such as its ability to bind/localize to curved microtubule protofilaments/plus-ends and diffuse on microtubules, mirrors the Ska1 complex. Further, both the Ska1 complex and Cdt1 carries a winged-helix domain that constitute their microtubule-binding domain [6, 10]. Finally, there is evidence that the Ska1 complex might require the loop domain to attach to the Ndc80 complex, like Cdt1 [11]. Thus, we conjectured that along with binding to microtubules directly, Cdt1 might also be able to access microtubules indirectly through its interaction with the Ska1 complex. Our previous studies were able to successfully demonstrate a modest interaction between Cdt1 and Ndc80 both in cells and *in vitro* [3, 6]; however, whether Cdt1 and the Ska1 complex would interact, remained unknown. To understand further how the formation of load-bearing k-MT attachments are generated and stabilized, we put-forth two key hypotheses; (a) The Ska1 complex and Cdt1 binds microtubules synergistically and (b) Cdt1, Ska1 complex and Ndc80 interact with each other to generate a trimolecular complex that is important for k-MT coupling.

In this study, we use a combination of biochemical, biophysical, and cell biological approaches to investigate these outstanding questions. Prior work has identified the Ndc80 as the major microtubule-binding complex at the kinetochore, however the complex has demonstrated no strong preference to bind to the growing or shortening MT tips [9, 12]. Although Ndc80 has been shown to be able to weakly track dynamic ends in combination with Ska1, the molecular milieu of a robust tip tracker during later stages of mitosis is still missing. Here we show that Cdt1 synergistically interacts with Ska1 and Ndc80 and together they form a tripartite complex that robustly tracks the depolymerizing end of dynamic microtubules.

## Results

### An auxin-induced degron cellular system to assay for mitotic functions of Cdt1

We had previously demonstrated the novel mitotic function of the replication licensing protein, Cdt1, using a double-thymidine cell synchronization approach coupled with RNAi or by injecting a function-blocking antibody into mitotic cells. Using this conventional approach, Cdt1 was degraded in the S-phase of the cell cycle and cells entering G2/M in the presence of Cdt1-siRNA were devoid of the newly accumulating Cdt1 during mitosis. However, it is still possible that these methods were not optimal in inhibiting the mitotic functions of Cdt1 and/or were also detrimental to the cells. We hence developed a new Auxin-induced Degron (AID) system for Cdt1 that drives its degradation upon Auxin, IAA addition. Both the genomic Cdt1 loci were replaced with a YFP- and degron-tagged version of Cdt1 in the resulting DLD1 stable cell line, hereafter referred to as AID-Cdt1 cell line (Fig. 1A, Sup. Fig. 1A). Using this approach, we anticipated that any cell entering mitosis after induction with IAA would not have encountered Cdt1 inhibition in any previous stage of its cell cycle outside of the M-phase.

**Figure 1:**
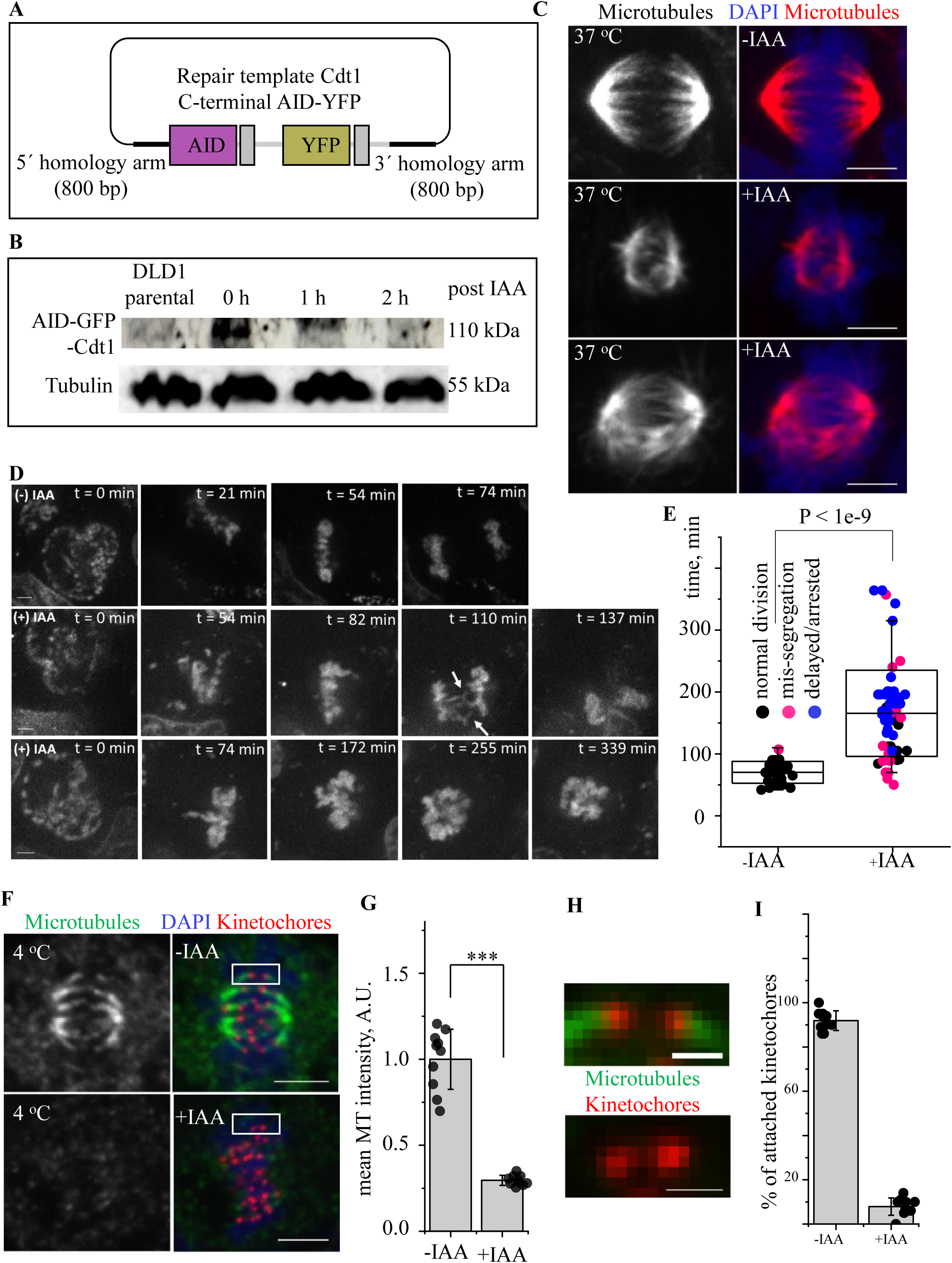
Generation of an auxin-induced degron (AID) cellular system to assay for mitotic functions of Cdt1 (AID-Cdt1). (A) A schematic of the repair template generated to endogenously tag the Cdt1 genomic locus with YFP and AID using CRISPER/CAS9 based genome editing. (B) Western-blot analysis showing the levels of the tagged Cdt1 in parental and AID-Cdt1 clones following auxin mediated degradation. (C) AID-Cdt1 DLD1 cells growing at 37°C were treated with either control DMSO (top panel) or the Auxin, IAA, for 2 hr. (middle and bottom panel) followed by immunostaining using an antibody against tubulin and the chromosomes counterstained with DAPI. Scale Bar, 5 µm. (D) AID-Cdt1 DLD1 cells growing at 37°C were treated with the DNA stain, Hoechst, along with either control DMSO (top panels) or IAA, for 2 hr. (middle and bottom panels) followed by live imaging for 2-6 hr. as required. Selected frames from the time series are shown for each experimental condition as indicated. Scale Bar, 5 µm. (E) Quantification of live imaging data from D. n = 30 for control cells not treated with IAA and 50 for cells treated with IAA. (F) AID-Cdt1 DLD1 cells growing at 37°C were treated with either control DMSO (top panel) or IAA, for 2 hr. (bottom panel) followed by treatment with ice-cold buffer PBS buffer for 10 min. The cells were then immunostaining for antibodies against tubulin, Zwint1 (for kinetochores) with the chromosomes counterstained with DAPI. Scale Bar, 5 µm. (G) Quantification of total microtubule intensities from cells in F (n=10 cells). (H) Insets from F showing samples of attached and unattached kinetochores from the top and bottom panels respectively. Scale Bar, 1 µm. (I) Quantification of the % of attached kinetochores from F under the two experimental conditions assessed (n=10 cells).

Efficient Cdt1 degradation was observed within 1-2 h of induction with IAA (Fig. 1B). However, to validate the AID system, we wanted to confirm if we could recapitulate the phenotypes which were previously observed using the traditional approach. We had demonstrated that Cdt1 depletion in mitotic cells led to substantial disassembly and destabilization of kMT attachments after cold treatment. Exposing the AID-Cdt1 cell line to IAA, we observed a similar phenotype. While at 30 min post-IAA exposure, there were still a considerable number of cold-stable microtubules retained in the metaphase cells, most of these microtubules completely disappeared within 1-2 h (Sup. Fig. 1B). Interestingly, unlike the siRNA-based depletion, a substantial fraction (∼ 80-90 %) of mitotic cells had severe defects in the structure of the mitotic spindle even at 37 ^°^C. Among these, majority of the cells (∼80%) had a disorganized metaphase spindle while, in a smaller fraction (∼ 20%), the spindles appeared rudimentary in size. Also all the cells in both these categories invariably exhibited substantial chromosome misalignment (Fig. 1C). Moreover, in all cells that were in anaphase or telophase exhibited at least one chromosome mis-segregation event (Sup. Fig. 1C.

To better characterize the phenotypes of AID-Cdt1 cells during mitosis, we carried out live imaging experiments of cells entering mitosis, after labelling them with the cell permeable DNA dye, Hoechst. The live imaging was initiated at the point of nuclear envelope breakdown (NEBD) and continued for a period of 2-6 h. While the uninduced control cells (AID-YFP-Cdt1) treated with either DMSO or Hoechst were YFP-positive prior to the initiation of imaging, the cells treated with IAA were YFP-negative. The control cells that were not subjected to Cdt1 degradation segregated their chromosomes in an average of 70 min after NEBD (n = 30; Fig. 1D top panel, Fig. 1E). However, the cells subjected to induced degradation of Cdt1 showed two broad categories of phenotypes, barring the very rare cell that underwent normal chromosome segregation (n = 55). The 1^st^ category constituted mitotic cells that exhibited mild to moderate or sometimes severe delays in mitotic progression and underwent chromosome mis-segregation at anaphase onset (Fig. 1D, middle panel; Fig. 1E). The 2^nd^ category of cells remained arrested in the mitotic state for the entire period of imaging with severe chromosome misalignment (Fig. 1D, bottom panel; Fig. 1E). Overall, the AID-YFP-Cdt1 cells spent extensively longer time (∼ 2.4 fold, 166 min) in mitosis, before a few cells proceeding to erroneous anaphase (Fig. 1E). Both the control cells and the parental DLD1 cells, as well as the cells devoid of Cdt1, rarely exhibited multipolar (tri- or tetrapolar) mitosis and these cells were excluded from our phenotypic analysis (not shown).

Next, we wanted to assess the stability of k-MT attachments in AID-Cdt1 cells after Cdt1 degradation. We observed a 75% loss of total microtubule fluorescence in Cdt1-depleted cells as compared to the control (Figs. 1F, G). The kinetochores in the AID-Cdt1 cells with IAA treatment were less stretched or tensed reflecting an absence of load-bearing attachments, in comparison to the control cells. Further, we also evaluated the frequency of physical contacts made between kinetochores and microtubules in both control and Cdt1-degraded cells. After Cdt1 was degraded, only ∼ 10% of the kinetochores showed proper bi-oriented attachments with spindle microtubules in contrast to the control cells, where ∼90% of the kinetochores were properly attached and bi-oriented (Figs, 1F, insets, Fig, 1H).

### Depletion of the Ska1 complex abrogates the localization of Cdt1 to kinetochores and spindle microtubules

Besides Cdt1, many proteins/protein complexes are recruited at the k-MT interface by the Ndc80 complex to facilitate stable k-MT attachments. One such kinetochore-localized complex of prominence is the metazoan Ska1 complex [8, 11, 13, 14]. Ska1 is a hetero-hexameric complex, composed of two copies each of SKA1, 2, and 3 subunits. Like Cdt1, the ability of Ska1 complex to dock on to kinetochore-bound Ndc80 and bind to microtubules to generate robust k-MT attachments prompted us to evaluate if the Ska1 complex shares a functional relationship with Cdt1. Moreover, as described earlier, in our search for proteins that could potentially boost the ability of Cdt1 to bind to microtubules, Ska1 emerged as the most obvious and attractive candidate worth testing. How these MAPs coordinate with Ndc80 either alone or in combination to modulate the strength of k-MT attachments in a stage-specific manner during mitosis was unexplored.

Our previous studies suggested that Ska1 levels at the kinetochores remained unaffected in the absence of Cdt1 [3]; however it is not clear if the Ska1 complex has a role in Cdt1 localization to the kinetochores. To address this question, we carried-out immunostaining of endogenous Cdt1 in HeLa cells that were treated with either scrambled siRNA (control) or siRNA against SKA3. While control cells showed distinct localization of Cdt1 to both kinetochores and the spindle poles (similar to SKA3) where density of microtubules is the highest; Cdt1 failed to localize properly to either of these structures in SKA3-depleted cells (Fig. 2A). Quantification of Cdt1 and Ska1 levels at the kinetochores suggests a positive correlation between the loss of Ska3 and Cdt1. An efficient knockdown of Ska3 led to a substantial loss of Cdt1 at the kinetochores (Fig. 2B; Sup. Fig. 2B). However, in line with our previous findings, in the reverse experiment where mitotic cells were subjected to siRNA-mediated knockdown of Cdt1, the levels of Ska3 at the kinetochores were unaffected (Sup. Figs. 2A; 2C-E)[3]. Our results thus point towards a hierarchical recruitment of these proteins during mitosis where Cdt1 is dependent on both the Ndc80 and the Ska1 complexes for its localization to kinetochores, while Ska1 only depended on the Ndc80 complex.

**Figure 2:**
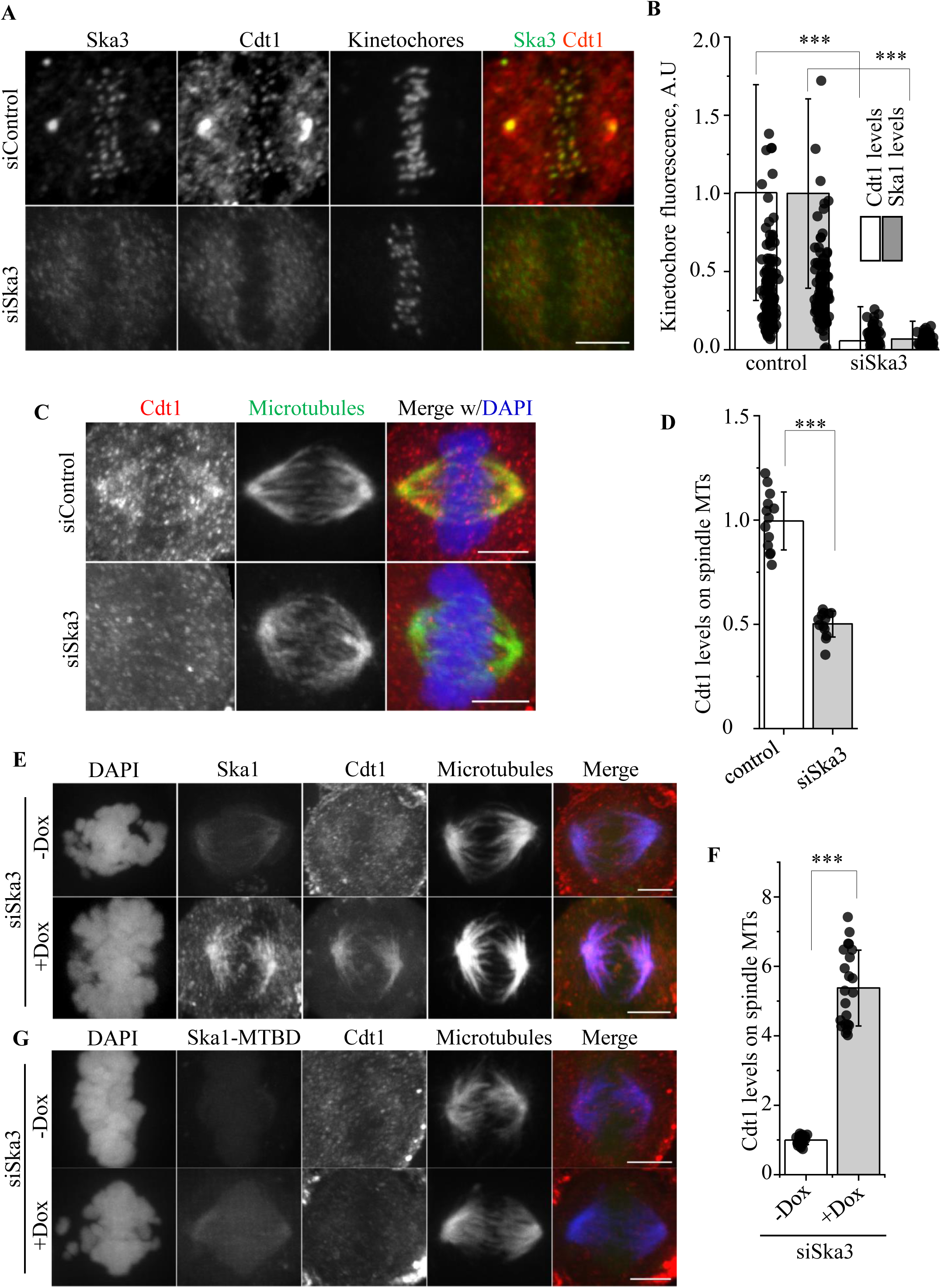
Ska1 complex is essential for the recruitment of Cdt1 to the kinetochores and spindle microtubules. (A) HeLa cells treated with either scramble control or siRNA against endogenous SKA3 were fixed using paraformaldehyde. Representative images of cells immunostained with antibodies against Hec1 (kinetochore marker) in far red, SKA3 in green and Cdt1 in red are shown. Scale bar, 5 μm. (B) A bar graph showing the quantification of kinetochore staining intensity after siRNA treatment in each case after background subtraction (n=110 kinetochores, > 5 cells). *p<0.05* represents statistical significance as assessed by the two-sided unpaired non-parametric student’s *t* test. (C) HeLa cells treated with either scramble control or siRNA against endogenous SKA3 were fixed using methanol. Representative images of cells immunostained with antibodies against Tubulin in green and Cdt1 in red are shown; DAPI stained chromosomes are shown in blue in the merged image. Scale bar, 5 μm. (D) A bar graph showing the quantification of Cdt1 spindle microtubule staining intensity after SKA3 siRNA treatment after background subtraction (n=13 cells). *p<0.05* represents statistical significance as assessed by the two-sided unpaired non-parametric student’s *t* test. (E-G) HeLa cells stably expressing either GFP-SKA1 (E and F) or GFP-SKA1^ΔMTBD^ (G) were treated with SKA3 siRNA to knockdown the endogenous SKA3 and the expression of siRNA resistant GFP-SKA1 or GFP-SKA1^ΔMTBD^ proteins were induced by adding 2.5 µg/ml of Doxycycline (+Dox) for 48 hr. Control cells were treated similarly with the siRNA in each case but no doxycycline was added (-Dox). The cells were then fixed and immunostained with antibodies against GFP for SKA1-FL or SKA1 ^ΔMTBD^ (2^nd^ column), Cdt1 (3^rd^ column and in red in the merge images) and microtubules (4^th^ column and in blue in the merge images) with the chromosomes counterstained using DAPI (1^st^ column). Scale bar, 5 μm. (F) A bar graph showing the quantification of Cdt1 spindle microtubule staining intensity from E in the presence and absence of Doxycline induction (n=22 cells). *p<0.05* represents statistical significance as assessed by the two-sided unpaired non-parametric student’s *t* test.

The next question was to test whether the binding of Cdt1 to microtubules was also dependent on the Ska1 complex, like its kinetochore recruitment. To answer this question, HeLa cells were immunostained to discern the level of endogenous Cdt1 on microtubules in the absence of the Ska1 complex. Indeed, the cells treated with SKA3 siRNA demonstrated ∼ 55 % reduction in the Cdt1 localization on the spindle microtubules (Figs. 2C, 2D) as compared to the controls, where Cdt1 clearly co-localized with the spindle microtubules in accordance with our previous study [6]. However, it is noteworthy that the overall spindle structure was also disrupted to a certain extent in the absence of the Ska1 complex (data not shown). Therefore, it was plausible that the inability of Cdt1 to localize to microtubules upon SKA3 knockdown could be attributed to the disruption of the spindle microtubules. To address this possibility, we performed a reverse knockdown-rescue experiment wherein, we expressed the siRNA resistant, microtubule-binding SKA1 subunit of the Ska1 complex in fusion with GFP in HeLa cells and asked whether Cdt1 localization on spindles could be enhanced. The siRNA resistant SKA1-GFP protein was induced by adding doxycycline and the endogenous SKA1 was depleted using siRNA. The over-expressed Ska1-GFP localized on the spindle microtubules as expected. More importantly, the confocal micrographs and the quantification of the fluorescence intensity signals clearly revealed an extensive and distinct localization of endogenous Cdt1 on spindle microtubules (Fig. 2E, 2F), which was around 5-fold higher than the uninduced control (-doxycycline) and also higher compared to the endogenous Cdt1 staining in normal HeLa cells (Fig. 2C) [6]. The results clearly demonstrate that the expressed SKA1-GFP was able to recruit substantial levels of Cdt1 to the mitotic spindle, pointing towards a possible mechanistic synergy between these two proteins for microtubule-binding. Interestingly, over-expression of the SKA1-GFP fragment (SKA1^ΔMTBD^, a. a. 1-131) devoid of the microtubule-binding domain (MTBD) was not able to recruit Cdt1 to microtubules as in the case of full-length SKA1-GFP (Fig. 2G). This could be attributed to the inability of the SKA1^ΔMTBD^ to localize to microtubules and/or the involvement of this domain in interacting with Cdt1.

Finally, to assess if Cdt1 depletion had any effect on the ability of Ska1 complex to interact with microtubules, we depleted Cdt1 from HeLa cells using Cdt1 siRNA coupled with double thymidine synchronization and evaluated endogenous Ska1 staining on microtubules. Our results demonstrated that upon the addition of Cdt1 siRNA, the localization of Ska1 on the spindle microtubules remained unaffected in comparison to the control untreated cells (Sup. 2C). The results clearly confirm that although Cdt1 is dependent on the Ska1 complex for both its kinetochore and microtubule localization, Ska1 recruitment as such did not require Cdt1.

### Cdt1 and the Ska1 complex interacts in cells and in vitro

The above data not only demonstrate the mechanism of how Cdt1 is recruited at the kinetochores via its interaction with the Ska1 complex; but also point strongly towards the possibility that these two proteins can also interact with each other. Therefore, we employed several biochemical methods, including GST pull down assay, co-immunoprecipitation (coIP), *in vitro* Blot overlay assay and Biolayer Interferometry (BLI) to validate our hypothesis.

For the GST pull down assay, mitotic HeLa cell lysate was incubated with equal amounts of either GST alone or GST-tagged Cdt1^92-546^ proteins (Fig. 3A). Upon elution with reduced glutathione, the glutathione agarose beads could pull down GST-tagged Cdt1 as expected, but SKA1 and SKA3 were also co-eluted and enriched in the bound (B) fractions. As control, GST was not able to precipitate the Ska1 complex as is evident from the detection of SKA1 and SKA3 only in the unbound (UB) fractions (Figs. 3A, 3B).

**Figure 3:**
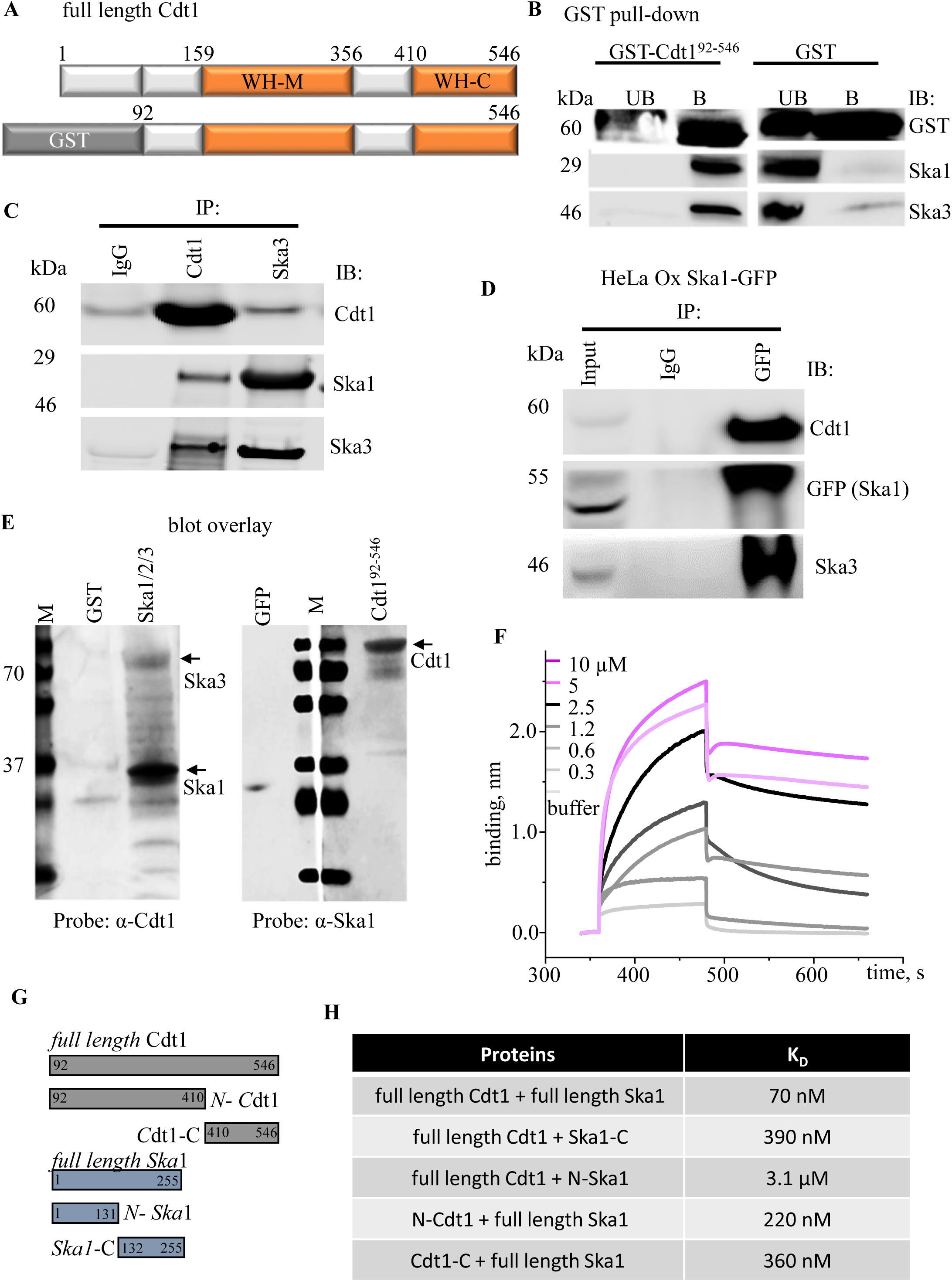
Cdt1 and the Ska1 complex physically interact with each other. (A) Diagrammatic representation of the Cdt1 construct purified from bacteria that were used for pull-down experiments. Structural domains relevant for the current study has been depicted. (B) Thymidine synchronized and nocodazole arrested mitotic HeLa cell extracts were incubated either with purified GST or GST-tagged Cdt1^92-546^ proteins (10 μg each), followed by pull down with glutathione agarose beads. UB indicates the unbound or flow through fraction and B indicates the proteins retrieved after elution with reduced glutathione. The blots were probed with anti-GST antibody and SKA1 and SKA3 antibodies. (C) Thymidine synchronized and nocodazole arrested mitotic HeLa cell extracts were immunoprecipitated (IP) and immunoblotted (IB) with the indicated antibodies; IgG was taken as a negative control. (D) Thymidine synchronized and nocodazole arrested mitotic HeLa cells that were stably expressing GFP-SKA1 (induced by adding 2.5 µg/ml of doxycycline) were immunoprecipitated (IP) and immunoblotted (IB) with the indicated antibodies; IgG was taken as a negative control. 1% of the lysate was loaded as input. (E) Blot overlay assay to study Cdt1-Ska1 complex interaction. Indicated proteins (0.5 µg each) were loaded as baits on 18% SDS-PAG, transferred to nitrocellulose membrane and blocked with 5% SM-TBST. Indicated proteins (1 µg) were overlaid as prey protein on the membrane for 12 h at 4 °C. The blot was washed and probed with indicated antibodies followed by chemiluminescence. Arrows depict the proteins of interest and molecular mass standards are shown in kDa. (F) Biolayer interferometry (BLItz) sensorgrams obtained using GFP-tagged Cdt1^92-546^-loaded amine-reactive biosensors and indicated concentrations (in µM) of Ska1 complex used as analysts to generate a series of sensorgrams, with orange bold dotted lines indicating the start of the binding (left) and dissociation (right) phases after the baseline. Binding curves were fit globally to a 1:1 binding model to yield equilibrium dissociation constant (K_D_), and association (ka) and dissociation (kd) rate constants, tabulated above the graph. (G) Linear diagram of the full-length and truncated proteins used to map the Cdt1-Ska1 interaction domains. (H) Table for K_D_ values estimated from the BLI sensograms obtained for the pairwise binding of the indicated purified protein constructs.

Further, to ascertain if Cdt1 and Ska1 complex interact *in vivo*, we performed coimmunoprecipitation (coIP) experiments in HeLa cells, first under endogenous conditions. Mitotic HeLa cell extracts were subjected to immunoprecipitation with non-specific IgG (as a control), or antibodies targeted against Cdt1 and SKA3. Cdt1 antibody was able to pull down endogenous Cdt1 protein as expected but simultaneously, it also immunoprecipitated both SKA1 and SKA3 proteins. Similarly, in a reverse pull down assay, the SKA3 antibody was also able to immunoprecipitate Cdt1 from the cell lysates along with SKA3 and SKA1 as expected (Fig. 3C). For assessing Cdt1 and Ska1 interaction under conditions where one protein is over-expressed; we transfected plasmid expressing HA/His-tagged Cdt1 in HEK293T cells followed by arresting the cells in mitosis through thymidine-synchronization and nocodazole treatment. Ni^+2^- NTA agarose-mediated affinity precipitation of HA/His-tagged Cdt1 led to the co-precipitation of both SKA1 and SKA3 as interacting partners (Sup. Fig. 3A). The coIP assay was also performed in HeLa cells that were stably expressing either GFP-SKA1 (under the doxycycline inducible promoter) or GFP alone. While anti-GFP antibody could precipitate GFP-SKA1 or GFP as expected, Cdt1 was co-precipitated only with the former and not the latter (Fig. 3D, Sup. Fig. 3B); indicating that Ska1 complex and Cdt1 indeed form a stable complex *in vivo*.

The results obtained so far from the pull-down experiments in cellular extracts suggested the possibility that these two proteins could certainly interact with each other but whether this interaction is direct and does not require assistance from any other extraneous protein(s) or cofactor (s) remained to be discerned. To assess this, *in vitro* blot overlay assay was carried out using purified recombinant proteins. In either case where the Ska1 complex assembled by combining equimolar His-tagged SKA1/2 (Sup. Fig. 3C) and GST-tagged SKA3 was used a bait and Cdt1^92-546^ was overlaid or vice versa, i.e., GFP-tagged Cdt1^92-546^ was used as a bait and Ska1 complex was overlaid, the proteins showed evidence of direct binding (Fig. 3E). GFP and GST proteins (used as negative controls for the tags) were not able to bind to the Ska1 complex or Cdt1 respectively, attesting to the specificity of the interaction.

Having demonstrated a direct interaction between Cdt1 and SKA1, we proceeded to compute the affinity of the interaction between Cdt1 and Ska1 complex. To accomplish this, we took advantage of biolayer interferometry (BLI), which exploits changes in the interference patterns when two biomolecules interact. Cdt1^92-546^ (0.2 mg/ml) was immobilized on the His-reactive sensor and increasing concentrations of Ska1 complex (0.3-10 µM) were used as analytes. The association and dissociation kinetics are monitored in each phase, resulting in a kinetic binding sensorgram from which the association and dissociation rate constants (*k_a_* and *k_d_*) and the equilibrium binding constant (*K_d_*) were determined. The *K_d_* for Cdt1 and Ska1 complex was determined to be 70 nM (Fig. 3F). As a negative control, BSA protein was used as an analyte and lack of binding confirmed the authenticity of the interaction. Further, using a combination of C- and N-terminal fragments of Cdt1 and Ska1, we identified the domains within Cdt1 and the Ska1 complex that were important for their mutual interaction (Figs. 3G, 3H; Sup. Fig. 3D). The *K_d_* of the C-terminal fragment of Ska1 binding to Cdt1, although considerably higher compared to full-length Ska1, was 390 nM. This is much lower than the N-terminal Ska1 fragment, the binding *K_d_* of which to Cdt1 was 3100 nM (Figs. 3G, H, Sup. Fig. 3E). This data indicates that the MTBD of the Ska1 subunit is primarily responsible for Cdt1-binding and for the recruitment of Cdt1 on to microtubules, in agreement with our results from Fig. 2 (Figs. 2E-G). In the context of Cdt1 binding to SKA1, both the N-terminal (92-410 aa, 220 nM) as well as the C-terminal (410-546 aa, 360 nM) regions of Cdt1 had comparable binding affinities to the full-length Ska1; albeit they were less efficient as compared to the full-length Cdt1 (Figs. 3G, 3H Sup. Fig. 3E). This suggest that as for Cdt1 microtubule-binding, both the central (WH-M) as well as the C-terminal (WH-C) Winged-Helix domains of Cdt1 (Fig. 3A) are required to bind to Ska1 efficiently. It is interesting to note that the same regions of Cdt1 (92-546 aa) were required for direct binding of Cdt1 to microtubules *in vitro* [6].

### Phosphorylation of Cdt1 by Cdk1 at G2/M transition impacts its ability to interact with the Ska1 complex

We had previously shown that the phosphorylation of Cdt1 by the mitotic kinase, Aurora B impacts the ability of Cdt1 to bind microtubules. Consequently, the expression of Aurora B phosphomimetic mutants of Cdt1 (Cdt1-10D) induced severe phenotypes, including the loss of k-MT attachment stability and delay in mitotic progression [6]. Based on our current results, it is possible that the phenotypes we observed with the mutants could at least partly be due to their inability of Cdt1 to interact with the Ska1 complex. To investigate this, we immunoprecipitated HA-tagged Cdt1-WT and Cdt1-10D from mitotic HeLa extracts using anti-HA antibody and assessed the interaction of both the WT and phosphomimetic proteins with the Ska1 complex. Interestingly, both Cdt1-WT and Cdt1-10D were able to pull down SKA1 and SKA3 equally efficiently (Sup. Fig. 4A), suggesting that the phosphorylation of Cdt1 by Aurora B kinase does not ablate Cdt1’s ability to interact with the Ska1 complex. Thus, unlike the direct-binding of Cdt1 to microtubules, the interaction between Cdt1 and Ska1 complex seems to be independent of Aurora B-mediated phosphoregulation.

Interestingly, Cdt1 is known to acquire phosphorylation by Cdk1 kinase at the G2/M transition [15]. There are at least 5 consensus Cdk1 phosphorylation sites located towards the C-terminal region of Cdt1 (Fig. 4A). Since addition of negative charges by phosphorylation is a common mechanism to control protein binding to microtubules, we first tested if Cdk1 phosphorylation, like Aurora B phosphorylation, is also instrumental in regulating Cdt1 microtubule-binding. We purified WT and Cdk1 phosphomimetic (2E3D) versions of Cdt1 in bacteria [6] (Sup. Fig. 4B) and tested their ability to bind to microtubules using a microtubule co-pelleting assay. We were unable to discern appreciable differences between Cdt1-WT and Cdt1-2E3D in their ability to bind microtubules directly (data not shown).

**Figure 4:**
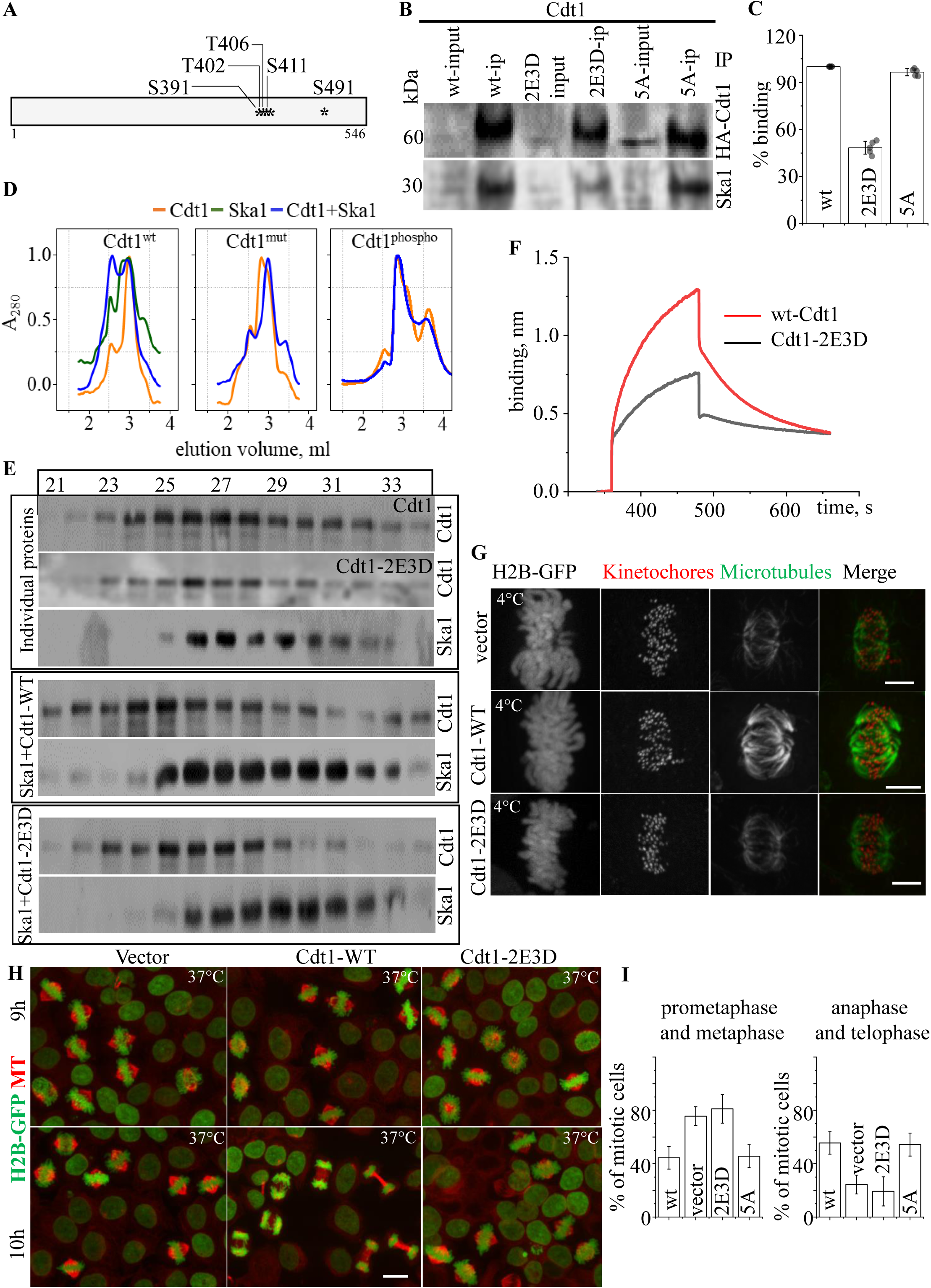
The phosphorylation of Cdt1 by the Cdk1 kinase interferes with Ska1-binding, impeding stable k-MT attachments and normal mitotic progression. (A) CDK1 Phosphorylation sites on Cdt1 that were mutated to obtain phophomemeitic and non-phosphorylatable versions of the same protein. (B)Western blot showing co-immunoprecipitation of HA-Cdt1 constructs and Ska1. (C) Quantitative band intensities from (B). (D) Elution profiles of indicated proteins or mixture of proteins from the co-fractionation experiments using Superose 6 gel filtration chromatography. (E) Western blots of fractions from (D) using primary antibodies as indicated on the right, for the proteins run separately or in combination as indicated on the left. (E) BLI sensogram of the indicated protein binding with Ska1. (G) Histone H2B-expressing, double thymidine synchronized, control HeLa cells (top panel) or those stably expressing RNAi-resistant wt (middle panel) or 2E3D mutant version (bottom panel) of Cdt1 were depleted of endogenous Cdt1 and the stability of K-fibers was assessed in each case by treatment with cold buffer 9 hrs after release from 2^nd^ thymidine arrest, as indicated. The cells were immunostained using anti-tubulin (green) and anti-Zwint1 (kinetochores, red) with the chromosomes counterstained using DAPI. (H) Same as in G, but the cells were fixed at both 9 and 10 hr. after release from 2^nd^ thymidine treatment followed by immunostaining using anti-tubulin (red) antibody. (I) Quantification of cells in various stages of mitosis from H.

Next, we generated stable HeLa cell lines expressing RNAi resistant versions of Cdt1-WT, non-phosphorylable (5A) and phosphomimetic (2E3D). We aimed to perform immunoprecipitation experiments using the cell lysates from these stable lines and test if these mutant versions were defective in binding to either the Ska1 or the Ndc80 complexes. Remarkably, we observed that both the WT and the 5A mutants of Cdt1 immunoprecipitated Ska1 equally well, but there was a 50-55% reduction in Ska1 within the immunoprecipitates of Cdt1-2E3D cell lysates (Figs. 4B, 4C). To further confirm this observation, we purified recombinant proteins, WT and 2E3D mutants of Cdt1 to perform co-fractionation experiments with the dimeric SKA1/SKA2 complex using analytical size exclusion chromatography. We used either Cdt1 (WT or 2E3D) or SKA1/2 separately or mixed them in equimolar ratio for 1 hr., before running them on the column. Both Cdt1-WT and -2E3D as well as SKA1 appeared to elute at their normal expected molecular weights when ran individually. However, when mixed, we observed a different elution pattern. A certain fraction of SKA1 appeared to coelute with Cdt1earlier than the expected elution point of this protein (Figs. 4D, 4E). On the other hand, we noticed that there was considerably reduced co-fractionation of SKA1 with the phosphomimietic Cdt1-2E3D (Figs. 4D, 4E). Similar results were also observed when a Cdk1-phosphorylated version of Cdt1-WT was used in this co-fractionation assay (Sup. Fig. 4C). Further, binding between SKA1 and Cdt1-2E3D protein was diminished as compared to the Cdt1-WT in the BLI experiment. The *K_d_* was reduced ∼2.5-fold for the mutant protein (70 nM *vs* 185 nM for the Cdt1-WT) (Figs. 3F, 4F, Sup. Fig. 4D). Together, our results suggest that the binding of Cdt1 and Ska1 is regulated by Cdk1-mediated phosphorylation.

Our attempts to test if the phosphorylation of Cdt1 by Cdk1 influences Cdt1’s ability to bind the Ndc80 complex yield results that was not in support of this notion. Our data indicated that both the WT, and Cdt1-2E3D were able to immunoprecipitate the Ndc80 complex to comparably similar extends (Sup. Fig. 4E).

### A Cdk1 phosphomutant of Cdt1 defective in Ska1-binding exhibits defective k-MT attachments and erroneous mitotic progression

We then sought to determine if the phosphorylation of Cdt1 by Cdk1 was physiologically relevant for mitotic cells. For this purpose, we utilized the stable HeLa cells lines expressing RNAi resistant WT, non-phosphorylable (5A) and phosphomimetic (2E3D) mutant versions of Cdt1 that we had used in our immunoprecipitation experiments with the Ndc80 and Ska1 complexes. As mentioned previously, we had derived an effective protocol to perform rescue experiments in double thymidine synchronized WT and mutant Cdt1 cell lines, where the endogenous Cdt1 has been degraded by siRNA-mediated knockdown [6](Sup. Fig 2A, 2B). The protocol exploits the fact that Cdt1 is degraded at the beginning of S phase and is freshly re-synthesized at the G2/M transition. When double thymidine arrested stable cells exogenously expressing RNAi-resistant Cdt1 is released from thymidine into the presence of Cdt1 siRNA, these cells enter mitosis 9-11 hr. after the release from thymidine with the newly synthesized, RNAi susceptible, endogenous Cdt1 being degraded immediately as it is being produced. We can now assess the mitotic functions of the exogenously expressed Cdt1 constructs.

The standard assay for Cdt1 mitotic function is the cold-stability assay as described previously (Fig. 1). Metaphase spindles of cells rescued by the WT or the non-phosphorylatable, Cdt1-5A were resistant to cold treatment as expected. On the other hand, spindles from the cells depleted of endogenous Cdt1 and rescued with the phosphomimetic, Cdt1-2E3D were highly susceptible to the cold treatment (Fig. 4G). We then monitored mitotic progression in these cells. Normally, HeLa cells released from thymidine arrest in S-phase will reach early mitotic prometaphase/metaphase, 9 h after release from thymidine, and after 10 hr., most of the mitotic cells in the synchronized culture will be in late mitotic anaphase/telophase. We observed that mitotic cells rescued by WT or the non-phosphorylable, Cdt1-5A mutant entered anaphase within the expected time, but the cells depleted of endogenous Cdt1 and rescued with the phosphomimetic, Cdt1-2E3D, remained in the early stages of mitosis even at the 10 h time point, suggesting that there was a delay in normal mitotic progression (Figs. 4H, 4I). Together, our results suggest that the perturbed Ska1-Cdt1 interaction in the cells expressing Cdk1 phosphomimetic (2E3D) mutant of Cdt1 resulted in defective k-MT attachments and delayed mitotic progression.

### Ska1 complex augments the ability of Cdt1 to bind to microtubules synergistically

Having determined that Ska1-binding is critical for Cdt1 targeting during mitosis and that this interaction is physiologically relevant, we next set out to reconstitute this interaction and test its relevance *in vitro*. In our recently published work, we have shown that Cdt1 can independently bind to microtubules with moderate affinity *in vitro* [6]. Since results from this work so far indicate a critical interaction between Cdt1 and Ska1 complex, we were tempted to postulate that affinity of Cdt1 to microtubules could potentially be enhanced by the addition of the Ska1 complex as was observed in the HeLa cells over-expressing SKA1-GFP. To validate this hypothesis, we employed TIRF microscopy (TIR-FM; Fig. 5A). For this purpose, we added sub-optimal amount of GFP-tagged Cdt1^92-546^ either alone or in combination with increasing concentrations of purified unlabeled His-Ska1/2 protein (Sup. Fig. 3C) on to the surface, immobilized with taxol-stabilized microtubules. While Cdt1^92-546^ alone showed only scarce and intermittent binding events at 1 nM concentration (Fig. 5B), it begins to start decorating microtubule lattice as the concentration of the SKA1/2 complex was increased. At 100 nM SKA1/2 complex, GFP-Cdt1 completely decorated microtubules and beyond that we observed a saturation in Cdt1 microtubule-binding (Figs. 5B, 5C). We also noticed a similar enhancement leading to the saturation of purified SKA1/2-GFP binding on microtubules with increasing concentration of untagged full-length Cdt1. This observation is confirmed by the quantification of intensities of Cdt1- and Ska1-GFP along microtubule lattices (Fig. 5D).

**Figure 5:**
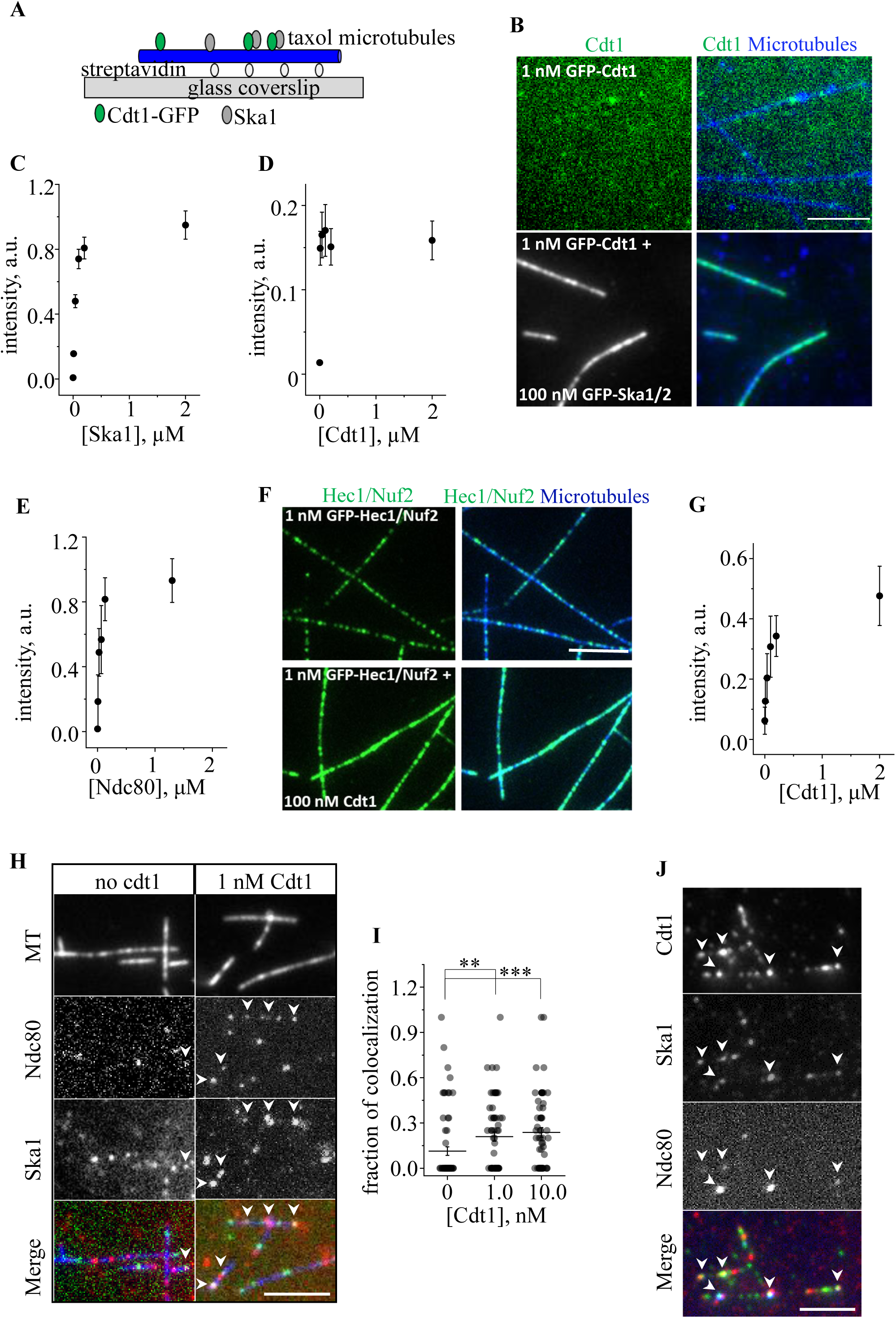
Cdt1, Ska1 and the Ndc80 complex exhibit synergy in binding to microtubules *in vitro*. (A) Schematics representation of the single molecule binding assay. (B) Selected images showing the binding of GFP-Cdt1 (1nM) on the Alexa 647-biotin-labeled microtubules either alone (top) or in presence of 100 nM untagged Ska1 (bottom). Scale bar 5 μm. (C) Data showing synergic binding experiment of GFP-Cdt1 (1 nM) with varying concentrations of untagged Ska1 (10 nM-2 µM). (D) is same as C but enrichment of GFP-Ska1 (1 nM) with varying concentrations of untagged Cdt1 (10 nM-2 µM) was plotted in this case. N≥2 experiments, n≥30 microtubules for each point, data are mean ± SEM. (E) Selected images showing the binding of GFP-Hec1/Nuf2 (1nM) on the Alexa 647-biotin-labeled microtubules either alone (top) or in presence of 100 nM untagged full length Cdt1 (bottom). Scale bar 5 μm. (F) Same as in C but showing quantification of synergistic binding between GFP-Cdt1 (1nM) and varying concentrations of GFP-Hec1/Nuf2. (G) Same as in D but showing quantification of synergistic binding between GFP-Hec1/Nuf2 (1nM) and varying concentrations untagged Cdt1. N≥2 experiments, n≥30 microtubules for each point, data are mean ± SEM. (H) Selected images showing binding at single molecule level of GFP-Hec1/Nuf2, and Ska1^Cy3^ proteins on taxol stabilized Alexa 647-biotin-labeled microtubules both in absence (left) and in presence (right) of 1 nM untagged full length Cdt1. Colocalized bindings are indicated with white arrowheads. Scale bar, 5 μm. (I) Scatter plot of the fraction of colocalized binding sites at three different concentrations of untagged Cdt1. Bar and whiskers are mean ± SEM. *** p<0.0001, ** p<0.001 (Mann-Whitney U-test). (J) Selected images showing binding at single molecule level of GFP-Cdt1, Ska1^Cy3^, and Ndc80^Alexa647^ proteins on taxol stabilized biotin-labeled microtubules. Colocalized bindings are indicated with white arrowheads. Scale bar, 5 μm.

Our previous work suggests a direct interaction between Cdt1 and the Ndc80 complex [3, 6]. Having observed the synergistic behavior between Cdt1 and SKA1/2 complex for microtubule-binding, we next sought to test if we could discern a similar synergy between Cdt1 and Ndc80 for microtubule-binding, using a similar TIR-FM approach used to analyze synergy with SKA1/2 complex. To accomplish this, we added sub-optimal amount of GFP-tagged Cdt1^92-546^ either alone or in combination with increasing concentrations of purified unlabeled Hec1/Nuf2 dimer of the Ndc80 complex on microtubules. While GFP-Cdt1^92-546^ at 1 nM concentration showed limited or no microtubule binding events, we observed that it started decorating the microtubules at 100 nM Hec1/Nuf2 dimer concentration, and attained saturation at even higher concentrations (Fig. 5E). More importantly, we observed a similar enhancement leading to the saturation of Hec1/Nuf2-GFP binding on microtubules with increasing concentrations of untagged full-length Cdt1 (Figs. 5E, 5G). These results suggest that Cdt1 not only exhibits synergy with the Ska1 complex for microtubule-binding but also with the Ndc80 complex.

The two-way synergistic binding of Cdt1 with both Ndc80 and Ska1 prompted us to hypothesize that Cdt1 can possibly form a tripartite complex with Ndc80 and the Ska1 complex once it is recruited to the kinetochores and microtubules and act as a co-factor to enhance the association between these two complexes. To test this, we resorted to single molecule TIRF imaging using sub nanomolar concentrations of fluorescently tagged SKA1/2^Cy3^,GFP-Hec1/Nuf2 dimer of the Ndc80 complex and untagged Cdt1^92-456^. The rational was that the presence of Cdt1 would increase the colocalization between the other two proteins. Indeed, we observed co-localized dots of GFP/Cy3 representing direct binding between GFP-Hec1/Nuf2 and Ska1^Cy3^ (Fig. 5H-left panel). Interestingly, the fraction of GFP/Cy3 colocalized dots doubled in the presence of Cdt1 at concentrations as low as 1 nM (Fig. 5H right panel). The fraction of colocalization increased upon increasing the concentration of Cdt1 up to 10 nM (Fig. 5I) beyond which the microtubule lattice was completely decorated with both Ska1^Cy3^ and GFP-Hec1/Nuf2 dimer and individual spots were not discernible. Furthermore, we also observed co-localization of all the three components (in ∼30% of cases) using sub nanomolar concentration of fluorescently tagged proteins (GFP-Cdt1, Ska1^Cy3^ and full length Ndc80^Alexa 647^) (Fig. 5J).

### The tripartite Cdt1-Ska1-Ndc80 super-complex tracks the ends of dynamic microtubules

While we have made substantial progress in obtaining mechanistic details about how robust k-MT attachments are formed and maintained during later stages of mitosis, we still have only very limited understanding of how kinetochores are stably coupled to dynamic microtubule plus-ends during chromosome alignment and segregation. Among the three proteins in context, Ska1 has been shown to be a weak tracker of dynamic microtubules, and although single Ndc80 molecules are not known to be a dynamic end tracker in metazoan systems, a combination of Ska1 and Ndc80 have previously been shown to be a weak tracker of microtubule ends *in vitro* [9]. Therefore, we investigated whether Cdt1 can impart more strength to these tip tracking complexes, Ndc80 and Ska1. For these experiments, we switched to polymerizable form of GMPCPP stabilized microtubule seeds [16] which can exhibit microtubule dynamics in the presence of soluble tubulin and Mg-GTP. Using the surface immobilized GMPCPP seeds (Fig. 6A), we observed normal binding of GFP-Cdt1 (1-100 nM) on the microtubule lattice. However, in presence of 1 mg/ml tubulin and 1 mM GTP, we did not observe any tip tracking activity of GFP-Cdt1 at any concentration tested (data not shown), indicating that Cdt1 might not be a bona-fide microtubule tip tracker. However, when GFP-Cdt1 (1 nM) was supplemented with 1 nM untagged SKA1/2 complex, weak tip tracking complexes were formed (4 % of the total number of microtubule seeds present in the field) (Fig. 6B, Sup Video 4). These complexes appeared to form in the solution and occasionally land near the ends of dynamic microtubules. A quarter of these complexes weakly track either the depolymerizing (20%) or polymerizing (5%) phase of the dynamic ends briefly. Less than 1% of these complexes found to follow both the polymerizing and the depolymerizing microtubule tips. Rest of these complexes were found to diffuse near the tips (Fig. 6D-left panel, and Sup. Fig. 6B).

**Figure 6:**
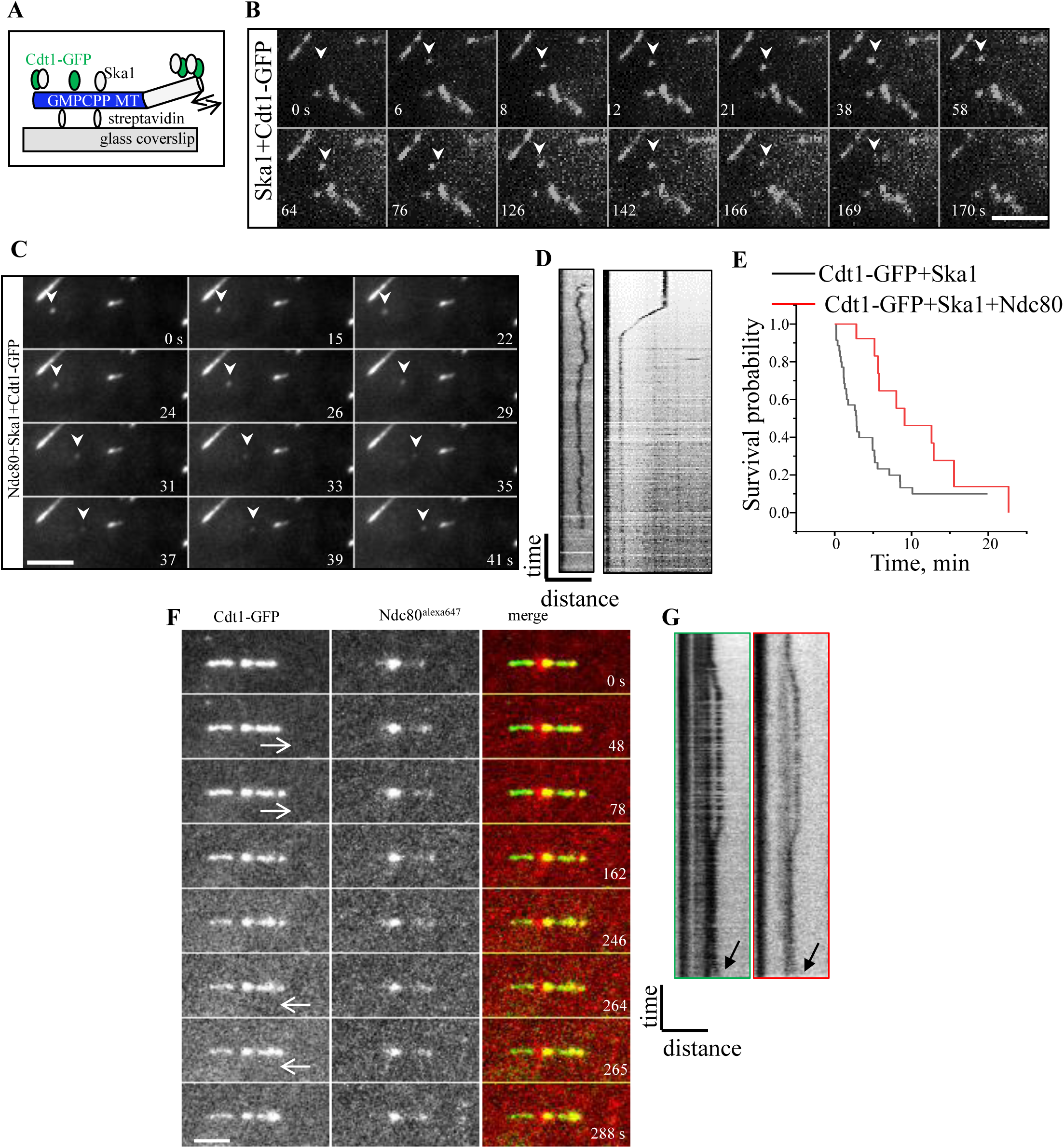
A tripartite complex comprising of Ndc80, Cdt1 and Ska1 exhibit potent tracking of depolymerizing microtubule plus-ends. (A) Schematic representation of a single molecule dynamic microtubule binding assay. Arrow showing the direction of microtubule dynamics. (B) Representative time lapse images recorded with surface immobilized microtubules after addition of a mixture of GFP-Cdt1 (1 nM), untagged Ska1 (1 nM), soluble tubulin (1 mg/ml), and Mg-GTP (1 mM), a complex of GFP-Cdt1 and Ska1 which lands near the tip is indicated with arrowheads. Time in s and the scale bar is 5 µm. (C) Same as in B but in presence of a mixture of GFP-Cdt1, untagged full length Ndc80 and untagged Ska1 (1 nM each). (D) Kymographs of the tip tracking complex in A (left), and B (right). (E) Kaplan-Meier survival curves for the total residence time (s) of tip tracking complexes formed in presence of indicated conditions. Calculated from n>30 events and N>3 independent trials in each case. (F) Representative time lapse images recorded with surface immobilized microtubules after addition of a mixture of GFP-Cdt1, Ska1, Ndc80^Alexa 647^ (1 nM each), and soluble tubulin (1 mg/ml), supplemented with Mg-GTP (1 mM). Arrow indicating the direction of movement of the tip tracking complex. Time in s and the scale bar is 5 µm. (G) Kymographs of the tip tracking complex in GFP-Cdt1 channel and in the Ndc80^alexa 647^ channel. Bar is 5 µm × 1 min. Arrow indicating regrowth event.

Surprisingly, when we add untagged Ndc80 (1 nM) together with GFP-Cdt1 (1 nM) and untagged SKA1/2 (1 nM) in this dynamic assay, we found similar landing of complexes on microtubule seeds (∼5% of the of the total number of microtubule seeds present in the field), but strikingly these complexes demonstrated far better tracking (Fig. 6C, Sup video 6) of the depolymerizing microtubule ends as evident form the kymographs (Fig. 6D, and Sup. Fig 6B). More than half (57%) of these complexes were found to be tracking either the polymerizing (21%), depolymerizing (21%) or both (15%) phases of microtubule dynamics. Most of these tip tracking complexes detached form the microtubule lattice after tracking the tip or diffusing near the tip, but some of them outlived our entire observation time of 30 min. To compare the residence time of these complexes and the differences between the bi/tripartite complexes, we computed survival curves, which showed that 50% of the tripartite (Cdt1-Ndc80-Ska1) complex survived for more than 8 min. whereas, only 20% of the bipartite (Cdt1-Ska1) complexes survived longer than 8 min. (Fig. 6E). Ndc80 has been previously shown to track the ends of dynamic microtubules together with Ska1 [9, 17], and in our hand, we also observed the formation of weak tip associated complexes with Ndc80 and Ska1 (Sup. Fig. 6A, Sup. Video 5). So, the enhanced tip tracking as observed with the tripartite complex could be ascribed to a possible upregulation of Ndc80-Ska1 binding mediated by Cdt1 and the consequent enhancement of end conversion of the tripartite complex. By using fluorescently labeled Ndc80^Alexa 647^, we also show that this tip-tracking tripartite complex (Sup. Video 7) actually contains both GFP-Cdt1 and the Ndc80 complex (Fig. 6F, G; Sup. Fig 6C). Furthermore, at least in a few instances, we have also observed tip tracking phenomenon lasting for more than one cycle of polymerization and depolymerization (Fig. 6G arrows). Finally, we also observed that the Ndc80 complex became more diffusive (Fig. 7A, 7B) as a part of the tripartie complex as compared to when used by itself with taxol-stabilized microtubules in our TIRF assays. Similar change in diffusivity was also observed for Cdt1 (Fig. 7C, 7D)

**Figure 7:**
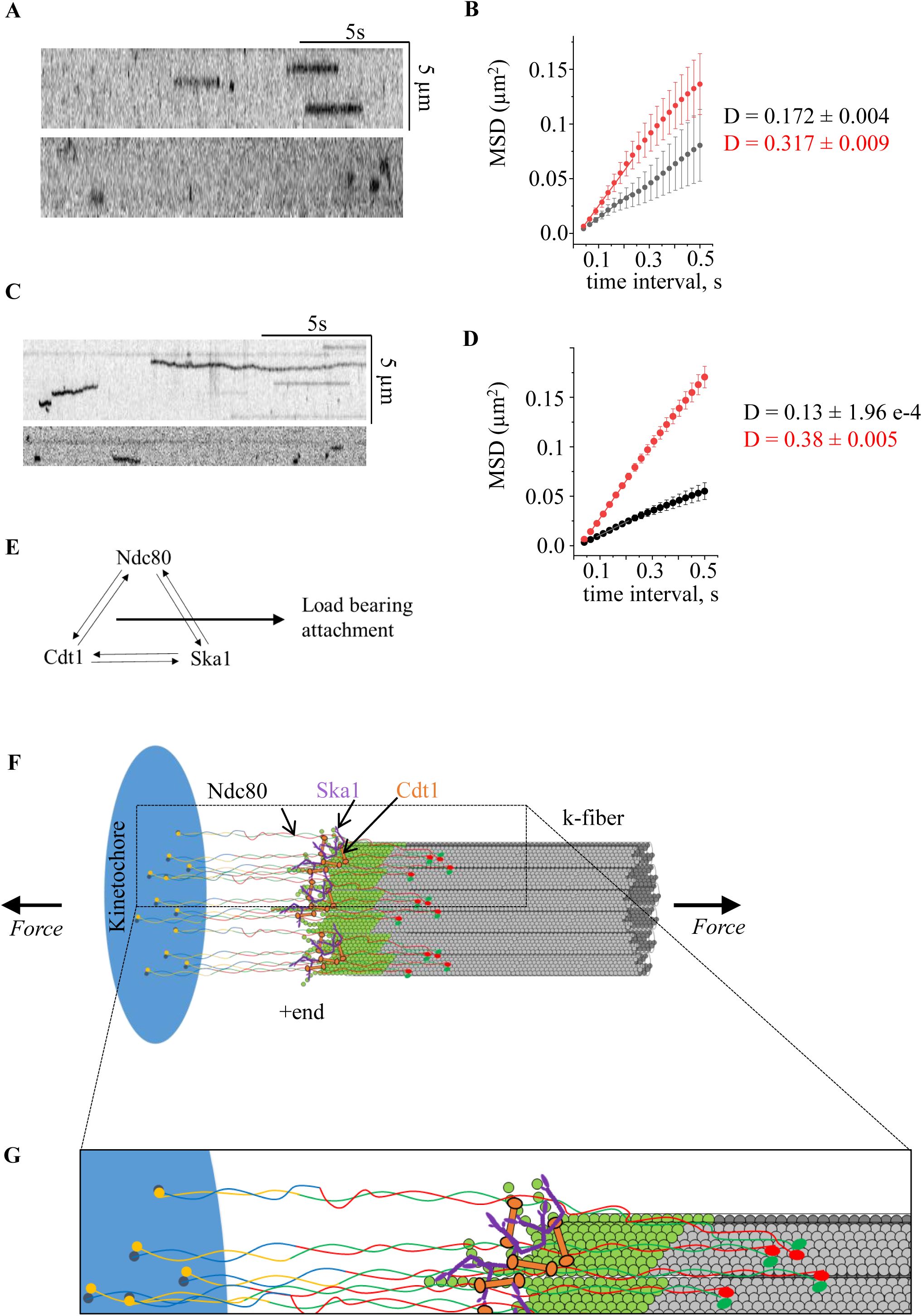
Processive end tracking activity to the tripartite Ndc80-Ska1-Cdt1 complex is characterized by enhanced diffusive behavior of the complex. (A) Single molecule diffusion kymograph for Ndc80-Alexa 647 alone (top), and Ndc80-Alexa 647 in the presence of GFP-Cdt1 and Ska1 (bottom). (B) Mean squared displacement plotted against time for Ndc80-Alexa 647 alone (black plot) and Ndc80-Alexa 647 in presence of GFP-Cdt1 and untagged-Ska1 (red plot). Straight lines representing linear fits to the data up to 0.22 s, the slope of which is the diffusion coefficient. (C) and (D) Same A and B but for GFP-Cdt1. (E) Mutual interactions between Ndc80 Cdt1 and Ska1 to form a super complex and the binding of the resulting super complex furnishes the load bearing attachment at kinetochore. (F) Schematics (not to scale) show multi-molecular lawn of Ndc80 molecules interacting with a kinetochore microtubule bundle. Propensity of Cdt1 (orange) and Ska1 (purple) proteins toward binding to curved microtubule protofilaments in conjunction with the intermolecular binding between Ndc80-Ska1-Cdt1 enables the formation of standalone microtubule tip tracking units. Green microtubules segments showing nascent GTP-tubulin cap. (G) Zoomed in section of k-microtubule attachment shown the Ndc80-Cdt1-Ska1 trio contacting the microtubules at two point of attachments one) on the microtubule lattice primarily by the CH-domains of Ndc80 proteins and second) Ska1-Cdt1 mediated interaction with the curved protofilaments flanked by Ndc80.

Overall, our data presents a novel finding that the Cdt1 interacts with Ska1 to facilitate a tripartite Ndc80-Ska1-Cdt1 complex that enables robust k-MT attachments and k-MT coupling for normal mitotic progression and accurate chromosomal segregation.

## Discussion

Faithful chromosome segregation is achieved by the co-operation between multiple molecular players and their regulatory factors. At the heart of this process are cellular mechanisms that facilitate the stabilization of k-MT attachments in metaphase and anaphase to prevent chromosome mis-segregation. The core microtubule-attachment machinery comprising primarily of the Ndc80 complex is well established to be critical for this stabilization. However, recent work has revealed that several accessory microtubule-binding proteins are also required for this process and more critically to efficiently couple the Ndc80 complex to the dynamic plus-ends of spindle microtubules [18, 19]. The Ska1 complex has been shown to be one such factor in several metazoan systems [8, 14, 20, 21]. Our work has demonstrated that the DNA replication licensing protein, Cdt1, is required for stabilizing k-MT attachments by facilitating robust connections between Ndc80 and spindle microtubules in humans [6, 22]. In this work, we demonstrate a viable interaction between Cdt1 and the Ska1 complex and provide evidence supporting the possibility that Cdt1 is an essential component of the k-MT plus-end coupling machinery during mitosis. Further, we show that Cdt1 enables k-MT coupling by synergizing with both the Ndc80 and Ska1 complexes to form a tripartite super-complex (Fig. 7A) that emerges as the most efficient tracker of dynamic microtubule ends to-date.

As detailed in the introduction, both the Ska1 complex and Cdt1 have been shown to directly bind to the Ndc80 complex. It is remarkable that Cdt1 and the Ska1 complex interact directly with a moderate to high affinity that is important to recruit Cdt1 to its mitotic targeting sites in the cells. The targeting of these proteins to kinetochore is hierarchical with the Ndc80 complex being most upstream and both Ska1 as well as Cdt1 being dependent on Ndc80. Cdt1 in turn is also dependent on Ska1, while Ska1 was found to be independent of Cdt1. This relationship between Ska1 and Cdt1 also seems to be true for spindle microtubule-binding in cells. However, interestingly, all these three factors can bind independently to taxol-stabilized microtubule *in vitro* while they also exhibit true synergy for microtubule-binding when used in pairs. The middle and C-terminal regions of Cdt1 comprising of the two winged-helix domains that are required for efficient Cdt1 microtubule-binding are also required for binding to Ska1 [18]. Further, the C-terminal winged-helix domain of Ska1 that is critical for its microtubule-binding is observed to be important for efficient Cdt1 spindle microtubule-binding in cells. Further, our studies find that Cdt1 associated more strongly with the SKA1 subunit than the SKA3 subunit of the Ska1 complex. Within the Ska1 complex, SKA1 functions as the primary microtubule binding subunit [9] and removal of its C-terminal domain abolishes the microtubule binding [23]. However, deleting the C-terminus of SKA3 only reduces the microtubule-binding affinity of the Ska1 complex by ∼ 4.5 folds [24]. We thus predict that Cdt1 most likely interacts primarily with the Ska1 subunit of the Ska1 complex. Finally, all three factors, Ndc80, Cdt1 and Ska1, when used in limiting concentrations show considerable colocalization on taxol-stabilized microtubules *in vitro*. Taken together, these observations point towards the existence of a tri-partite Ndc80-Ska1-Cdt1 complex in mitotic cells.

The k-MT end coupling-load bearing complex was long modeled upon the hill-sleeve model [25] in higher eukaryotes or the sliding ring model [26] for yeast, with other factors possibly assisting the Ndc80 complex in this process [18, 19. But considering that a Dam1-like ring complex is evolutionarily missing in metazoan eukaryotes and that their k-fibers are formed of 10-12 individual microtubules, direct experimental evidence in support of either of the models has not been established. Existing studies have not revealed a model where Ndc80 and Ska1 in concert can function via a ring or sleeve model. Our studies so far does not address the scenario whether Cdt1 can contribute to the formation of such a ring/sleeve around k-fibers consisting of multiple microtubules. Additional experimental approaches including Cryo-EM studies and force measurements using optical trapping reconstitution experiments are required to shed light on whether similar mechanisms exist in metazoan systems. With the existing evidence, we pose an alternate view that higher eukaryotes utilize a distinct k-MT coupling mechanism in which the three components, Ndc80, Ska1 and Cdt1 comprise single functional units of a processive end-tracking complex (Fig. 7B, 7C). This standalone unit, likely consisting of a single copy of each of the components can independently track the ends of dynamic microtubules. We have shown that this complex tracks not only the depolymerizing but also the polymerizing part of the dynamic microtubules, and rare instances we have also seen it tracking for multiple cycles (Fig. 6F-arrows). The fact that the observed tracking events were short-lived is likely a result of a missing a key kinetochore component such as kinesin CENP-E [in our assay] which is thought to maintain these end on attachments {Chakraborty, 2019 #286]. While both Ska1 and Cdt1 can bind curved protofilaments[6, 9], interestingly, only Ska1 seems to be able to transiently track plus-ends by itself and is required to target Cdt1 and Ndc80 to this site *in vitro*. Increased diffusion of Ndc80 as a part of the tripartite complex may be critical to impart microtubule plus-end tracking ability to the complex as has been predicted previously by our computational simulation experiments [27]. This observation is also supported by a concomitant increase in Cdt1 diffusion as a part of the complex. However, all three components comprising the standalone units are required for long distance processive tracking and Cdt1 seems to serve a critical function in this process.

Chromosome segregation is a very tightly controlled process, orchestrated by the interplay of multiple kinases/phosphatases. This is especially true for the regulation of k-MT attachments where multiple kinases (Aurora B, Cdk1 etc.) and phosphatases (PP1, PP2A, etc.) have been shown to regulate the interaction of factors at the k-MT interface with each other and/or with the spindle microtubules [28, 29]. The k-MT attachment strength also needs to be spatially and temporally regulated during different phases of mitosis. Our studies suggest that the replication licensing protein, Cdt1, is temporally regulated by Cdk1 to modulate the strength of k-MT attachment at different phases of mitosis. Cdk1 phosphorylation of Cdt1 is instrumental in regulating Ska1-Cdt1 interaction and for controlling the recruitment of Cdt1 to kinetochores and spindle microtubules. Previous work has demonstrated that Cdk1 activity is highest at G2/M transition and mitotic prometaphase and progressively reduces as the cells proceed to metaphase and anaphase [30]. Our results support a model where, at prometaphase Cdk1 phosphorylates Cdt1, reducing Cdt1 recruitment to kinetochores [27] and possibly accounting for weaker k-MT attachment earlier during mitosis. At metaphase, the kinetochores are correctly oriented, and the Cdk1 kinase levels are low, allowing for efficient Cdt1 recruitment via its interaction with the Ska1. Cdt1 and Ska1, in turn strengthen k-microtubule attachments in synchrony with Ndc80 and enable plus-end tracking ability to Cdt1-Ska1-Ndc80. Intriguingly, it is known that Cdt1 is not degraded at metaphase to anaphase transition and that Cdk1 phosphorylated Cdt1 is not competent for licensing the origins in G1 [15, 31]. This suggests that Cdt1 dephosphorylation would be critical for this cell cycle transition and thus future work is required to identify the phosphatase that contributes to this important function.

Future research is required to understand whether other molecules including chTOG, EB1, or Astrin-SKAP function as a part of this building block of if they function with the Ndc80 complex independently of this core unit. Also remains open is the question how this unit tracks a bundle of parallel microtubules [32] *in vitro* as opposed to single microtubules.

## Experimental procedures

### Cloning, recombinant protein expression and purification and labeling

GFP-tagged Cdt1 (92-546 a. a. referred through the text as GFP-Cdt1) and its subsequent deletion fragments (92-232, 92-410 and 410-546) were purified as described previously [6]. Plasmids encoding GST-tagged SKA3, GFP-His-SKA2, and His-tagged SKA1/2 for bacterial protein purification were kind gifts from Dr. Gary Gorbsky, A. Arockia Jeyaprakash and Dr. P. Todd Stukenberg laboratories respectively. The proteins were purified according to published protocols and aliquots were stored in -80 °C. Plasmids encoding His-SKA1/2 and GFP-His-SKA2 were transfected into BL21 *E. Coli* cells. Transformants were grown in 2xYT media at 37 °C until 1.0 Optical Density is reached. Protein expression was induced by adding 0.35 mM IPTG at 18 °C overnight. Cells were pelleted using centrifugation at 4412 *× g*. Pellets expressing SKA1/2 and GFP-His-SKA2 proteins were combined and the cells were lysed by sonication on ice in lysis buffer (20 mM Tris pH 8.0, 500 mM NaCl, 2 mM 𝛽-ME, supplemented with 1X protease Inhibitor (Thermo Scientific, #AA32965). Lysate was then clarified by centrifuge at 58387 × *g*, 4°C, for 30 min. Clarified supernatant was then incubated with Ni-NTA beads (Qiagen) at 4°C for 1 h on a roller. Flow through was discarded and, the beads were then washed with 30 column volume of high salt wash buffer (20 mM Tris pH 8.0, 1000 mM NaCl, 50 mM KCl, 10 mM MgCl_2_, 2 mM 𝛽-ME). The protein complex was eluted with elution buffer (20 mM Tris pH 8.0, 100 mM NaCl, 400 mM Imidazole, 2 mM 𝛽-ME). Fractions were evaluated with SDS PAGE electrophoresis, and the ones containing both His-SKA1/2+GFP-His-SKA2 subunits were pooled and further purified using size exclusion chromatography over a Superose 6 Increase 10/300 GL (GE Healthcare) column equilibrated with gel filtration buffer (20 mM Tris pH 8.0, 150 mM NaCl, 2 mM 𝛽-ME, 2% v/v glycerol). Fractions were analyzed again using SDS-PAGE and the ones containing both His-SKA1/2+GFP-His-SKA2 subunits were pooled and aliquoted. Aliquots were flash frozen and stored at -80 °C. Purified GFP-Hec1/Nuf2 dimer of the Ndc80 complex used for TIRF imaging was a gift from Arshad Desai. The whole human Ndc80 complex, also used for TIRF imaging was a gift from Andrea Masaccio laboratory and was labeled with Alexa 647 dye (Thermo Fisher Scientific, #A20006) using succinimide chemistry and following manufacturer’s protocol. Full length Ska1 complex was also labeled in an analogous way with Cy3 dye (Amersham, #PA13104). The labeling reaction produced homogeneously labeled fluorescent proteins of which the majority of them showed single step photobleaching patterns, as observed with GFP-Cdt1 (Sup. Fig. 5)

### Auxin-inducible degron tagging of endogenous Cdt1

We downloaded the human genomic sequence of Cdt1 from NCBI (NC_000016.10:88803778-88809258 Homo sapiens chromosome 16, GRCh38.p12) using SnapGene and located the exon/intron boundaries in this annotated sequence. We input the last exon sequence (+800 bp up/down stream) to design our guide RNA targeting the Cdt1 C-terminus, using the Broad Institute sgRNA design tool. We then cloned the gRNA into PX330 (obtained from Addgene #42230), a CAS9-EGFP sgRNA vector, following established protocols [33–35]. We confirmed the insertion by DNA sequencing. For the designing of the Cdt1 repair template, we synthesized a DNA fragment containing the above mentioned 5- and 3- 800 bp of Cdt1 homologous sequence with the sequence of the AID and YFP inserted in between these two homology arms (GeneUniversal). We then cloned this product into the vector HJURP C-terminal AID-YFP at KpnI-HindIII restriction sites (a kind gift from Dr. Daniel Foltz, Northwestern University). The PAM sequence within Cdt1 homology arms were mutated. DLD-1 Tir1 parental cells (a kind gift from Andrew Holland, Johns Hopkins University) were transfected with the plasmid containing the Cdt1 gRNA/Cas9 and the Cdt1 repair template at equimolar ratio using Effectene (Qiagen, #301425) transfection agent. Following transfection, the cells were grown for 7 days, after which they were collected by centrifugation at 200 × *g* for 5 min. The cells were then resuspended in 500 µl of 1% FBS in PBS. A 96 well plate was prepared with 100 µl of condition media collected from a confluent plate of the parental cell line after filtering using a 0.22 µm filter. We then sorted single cells into 96-well plates using Fluorescence Activated Cell Sorting at the Northwestern University – Flow Cytometry Core Facility. Then single cells were allowed to grow in the 96 well plate using complete DMEM media supplemented with 20% FBS. Single colonies were then transferred into 24-well plates, followed by 12-well and then finally to 6-well plates to ensure enough propagation of the single clones. To check the transfection efficiency, we first assessed YFP expression levels for each clone using confocal microscopy, then proceeded with making genomic DNA of each positive clone and identification of homozygous genomic insertion clones by PCR. The cloning strategy used yielded a 0.5 kb amplification for wild-type cells, a 1 kb amplification for homozygous insertions and bands of both size for heterozygous insertions (Sup. Fig. 1A). Monoclonal lines were further screened by Western Blotting and genotyping. We treated both control DLD1 cells and AID-DLD1 (clone#6) cells with 0.5 mM Indole Acetic Acid (IAA Auxin) at the following time points, 0, 1, and 2 hr. Cells induced with IAA for these different time points were lysed and analyzed for the levels of YFP-AID-Cdt1 with Western Blotting using an anti-GFP antibody.

### Immunofluorescence and live cell imaging

Methanol fixation was used for spindle microtubule staining for Cdt1 while Paraformaldehyde fixation was used for kinetochore staining. The rest of the details for immunofluorescence staining and confocal microscopy was exactly as described in our previous work [6]. The antibodies used for immunofluorescence include β-tubulin monoclonal (DM1A, Sigma), Zwint1 polyclonal (A300-781A, Bethyl Laboratories), Cdt1 polyclonal (Santa Cruz Biotech), Hec1 monoclonal (GTX70268, GeneTex) and anti-human ACA (Immunovision, HCT-0100). Guinea Pig Cdt1 antibody [3] was a gift from Jeanette Cook’s lab while SKA1 and SKA3 rabbit polyclonal antibodies were a gift from Gary Gorbsky.

Live imaging of chromosomes during mitosis in DLD1 cells were also carried out exactly as described previously for HeLa cells [6] using cell permeable DNA dye Hoechst. However, in this case, the DLD1 cells were not synchronized and the 35 mm glass-bottom MatTek dishes were manually coated with 10 µg/ml Fibronectin prior to using them for cell culture. Images of the Hoechst-labelled chromosomes were acquired every 7.5 minutes for up to 2-6 hrs as required based on whether the samples were controls or if Cdt1 was degraded.

### Analytical size exclusion chromatography

Untagged Ska1 complex and either of GFP-Cdt1 or GFP-Cdt1^2E3D^ proteins were mixed at concentrations 3 mg ml^-1^ and 1.6 mg ml^-1^ respectively and were incubated at 10°C for at least 2 h. The mixture was then clarified by centrifugation for 10 min at 16100 × *g* at 4°C. The mixture was analyzed by size exclusion chromatography using Superose 6 Increase 5/150 GL column (GE Healthcare) mounted on an AKTApure system. The column was equilibrated with size exclusion chromatography buffer (20 mM TRIS pH 8, 200 mM NaCl, 2% v/v glycerol, 2 mM TCEP) and the column was operated at a flow rate of 0.1 ml min^-1^. Typical fraction volume was 0.05 ml, the fractions were collected and analyzed by SDS PAGE electrophoresis and western blotting using anti-Cdt1 (Santa Cruz Biotechnology, # H-300, rabbit polyclonal IgG) and anti-SKA1 (Abcam, #ab91550, rabbit polyclonal) antibodies.

### Kinase assays

Purified GFP-Cdt1 protein (1.6 mg ml^-1^) was treated with CDK1-CyclinB Kinase (EMD Millipore, # 14-450) supplemented with 10 mM ATP in kinase buffer (20 mM Tris pH 8, 150 mM NaCl, 10 mM MgCl_2_, 1 mM DTT, 1X Phosphatase inhibitor (Thermo Scientific, #1862495)) at room temperature for 2 h. The phosphorylated GFP-Cdt1 protein was then incubated with Ska1 complex for 2 h at 10 °C. The mixture was then run on a size exclusion chromatography column (see above), fractions were collected and analyzed by SDS PAGE electrophoresis and western blotting as above.

### In vitro blot overlay assay

To assess interaction between the proteins of interest, 0.5 µg of one protein was loaded on to the SDS-PAG as “bait” and was transferred on to the nitrocellulose membrane using standard procedure. The membrane was blocked with 5% skimmed milk in TBS buffer and 0.1% Tween20 for an hour at RT followed by the addition of the second protein (1µg/ml in TBS) as “prey” for 18 h at 4 °C with constant shaking. The blot was thoroughly washed thrice for 15 min each with TBST and antibody to detect the prey protein, which was added at indicated dilutions for 2 h with constant shaking at RT. HRP-tagged mouse/rabbit antibodies (Jackson laboratories) were used at 1:10,000 dilution to detect the bait-prey complex and blot was developed by Super signal west pico chemiluminescent substrate (Thermofisher).

### Bio-layer Interferometry

Biolayer interferometry experiments were performed as in published protocols[36] on a single channel BLItz^®^ instrument (*forte*BIO, Menlo Park, CA). His-reactive biosensor tips (HIS1K, FORTEBIO, #185121) were hydrated in 96 well plates in BLI-buffer (20 mM HEPES pH 7.4, 1% sucrose, 150 mM NaCl, 0.5 mM TCEP, 0.02% Triton X-100) for 10 min, followed by ligand (0.2 mg/ml in BLI-buffer) immobilization for 300 s. Analyte in varying concentrations (0.3-10 μM) was placed as a drop (4 μl) onto the magnetic holder and was allowed to contact the sensor with the immobilized ligand for 120 s (association phase) followed by a dissociation phase for 180 s. Baselines for 30 s were obtained in BLI-buffer to compensate for the phase shifts. All the steps were performed at a constant shaking of 1000 rpm. Both the immobilized and analyte proteins were subjected to buffer exchange into the BLI buffer as required and pre-cleared using high speed ultracentrifugation to exclude the aggregates/debris prior to their use in BLI experiments. The temperature of measurement was set at 25°C unless otherwise specified. GFP and BSA proteins were taken as a negative control to rule out the possibility of non-specific interaction. Sensorgrams were fit globally to a 1:1 binding model by BLItz Pro version 1.1.0.28, from which the equilibrium dissociation constant (K_D_), association (k_a_) and dissociation (k_d_) rate constants, were calculated [36] (muller Esparza frontiers mol sci 2020).

### Cell culture, co-immunoprecipitation and western blotting

HeLa, HEK293T and GFP-expressing HeLa (kind gift from Dr. Robert Goldman, Northwestern University) cells were cultured at 37 °C with 5 % CO_2_ in Dulbecco’s modified eagle’s medium (DMEM, Life Technologies) containing 10 % fetal bovine serum (Seradigm, VWR Life Science), 100 U ml^−1^ penicillin and 1 μg ml^−1^streptomycin. HeLa stable cell lines expressing GFP-SKA1 clone D1 and GFP-SKA1^ΔMTBD (1-132)^ clone B6 were generously procured from Dr. Gary Gorbsky and were maintained similarly except for supplementing the medium with 150 µg ml^-1^ hygromycin.

The GFP-, SKA1-GFP and GFP-SKA1^ΔMTBD^ cell lines were grown till 50-60 % confluence. The cells were synchronized by treatment with 2 mM thymidine for 18 h followed by release for 9 h and then addition of 0.33 µM Nocodazole for 12 h to arrest the cells in mitosis. To induce the expression of GFP-SKA1 or GFP-SKA1^ΔMTBD^ (lacking the C-terminal residues, containing only 1-132 aa) proteins, doxycycline (2.5 µg/ml) was added in the medium from the time of synchronization. The mitotic cells were collected using mitotic-shake-off and were lysed in the co-IP buffer (10 mM HEPES, pH 7.4, 1.5 mM MgCl_2_, 10mM KCl, 0.1% Nonidet P-40 supplemented with 1 × Halt protease and phosphatase inhibitor cocktails, Thermo Fisher Scientific). One mg total cell protein was used per IP and 1% of the cell lysate volume was loaded as the input to assess the presence of desired proteins in each case. The antibodies (3 μg/IP) used include anti-GFP mouse monoclonal 3E6 (Thermo Fisher), Anti-GFP rabbit polyclonal A-11120 (Thermo Fisher) anti-SKA3 rabbit polyclonal ab186003 (Abcam), anti-Cdt1 rabbit polyclonal H-300 (Santa Cruz Biotech), or Control IgG. Magnetic Dynabeads^TM^ Protein G (Invitrogen) were used to isolate protein-antibody complex and were eluted by boiling the beads with 2 × Laemmli sample buffer (Bio Rad). Proteins were resolved on 4-15 % gradient gel and transferred using Western blot. The interacting partners were detected by Western blotting using relevant antibodies (Cdt1 H300, SKA1 and SKA2 polyclonal antibodies received from Dr. Gary Gorbsky and GFP A-11122, all at 1:1000 dilutions. TrueBlot-HRP antibodies against rabbit or mouse (1:1000) were used as secondary antibody (Rockland Immunochemicals Inc.). The blots were developed by chemiluminescence.

For co-IP experiments involving WT and Cdt1 phosphomutant variants (Cdt1^5A^, Cdt1^2E3D^, Cdt110^A^, and Cdt10^D^) and SKA1, stable HeLa cells expressing the respective proteins were grown till 70-80 % confluence, followed by the addition of 0.33 µM Nocodazole for 10-12 h to arrest the cells in mitosis. Cell lysate (1 mg per input) was incubated with mouse anti-HA magnetic beads (Thermo Scientific, #88836) at a ratio of 20:1 v/v. The interacting partners were detected by Western blotting using the antibodies: anti-HA rabbit polyclonal (Sigma, #H6908), and anti-SKA1 rabbit polyclonal (Abcam, #ab91550) at 1:1000 dilutions. For the co-IP of Hec1, rabbit anti-HA magnetic beads (Cell Signalling, #C29F4) was used for the pull-down and anti-Hec1 9G3 (GeneTex, #GTX70268) was used for Western blotting. HRP antibodies against rabbit and mouse (1:1000) were used as secondary antibodies (azure biosystems, #AC2114 and #AC2115, respectively) for detection and the blots were developed by chemiluminescence.

### Ni^+2^-NTA agarose-mediated pull down

HEK-293T cells were transfected with the plasmids (Cdt1-HASH in pQCXIP vector) for 6 h at 20-30% confluency using calcium chloride-based method. M phase synchronization was carried out by treating the cells with 2 mM thymidine for 18 h, followed by addition of 100 ng/ml (0.33 μM) Nocodazole for 10 h. The cells were collected by mitotic-shake-off and lysed using lysis buffer (50 mM HEPES pH 8.0, 33 mM KAc, 117 mM NaCl, 20 mM Imidazole, 0.1% triton, 10% glycerol, 0.1 mM AEBSF, 10 μg/ml pepstatin A, 10 μg/ml aprotinin, 10 μg/ml leupeptin, 1 mM ATP, 1 mM MgCl_2_, 5 μg/ml phosvitin, 1 mM β-glycerol-phosphate, 1 mM orthovanadate). The whole cell lysate expressing the His-tagged protein was incubated with Ni^+2^-NTA beads for overnight with end-on rotation at 4°C. Beads were washed three times with the lysis buffer, and bound proteins were eluted by boiling for 5 min in 40 μL of 2×SDS sample buffer. For immunoblotting, proteins were electrophoresed by SDS-PAGE and transferred to the PVDF membranes. Immunoblots were developed using chemiluminescence and exposed on to the X-ray films. Cdt1 H-300 polyclonal antibody (Santa Cruz) was used at 1:1000 dilution; SKA1 and SKA3 antibodies (1:1000 dilution each) were generous gifts from Dr. Gary Gorbsky.

### GST pull down assay

HeLa cells were synchronized at mitotic phase by adding 2 mM thymidine for 18 h followed by release for 9 h and subsequent addition of 0.33 μM Nocodazole for next 12 h. The mitotic cells were harvested by gentle shake-off and were washed twice with 1×PBS. The cell pellet was resuspended in the lysis buffer (10 mM HEPES, pH 7.4, 1.5 mM MgCl_2_, 10 mM KCl, 0.1 % Nonidet P-40 supplemented with 1 × Halt protease and phosphatase inhibitor cocktails, Thermo Fisher Scientific). The lysate was incubated with 10 μg of either GST or GST-tagged Cdt1^92-546^ proteins for 12 h at 4 °C with constant shaking. The lysate was then incubated with Glutathione resin 4B FF resin for 2 h followed by elution with 30 mM reduced glutathione in 50 mM Tris-HCl, pH 8.0. The proteins in the bound (B) and unbound (UB)/flow through were analyzed by Western blotting using anti-GST rabbit polyclonal antibody (Sigma #G7781, 1:5000 dilution).

### Total internal reflection fluorescence microscopy (TIR-FM)

#### Microtubule assembly

Taxol stabilized microtubules were prepared from mixing on ice the following; 4 µl 10 mg/ml unlabeled tubulin, 0.4 µl 5 mg/ml Alexa 647 labeled tubulin (Pursolutions #64705), 0.4 µl of 5 mg/ml biotin labeled tubulin (Cytoskeleton, #T333P-A), 20% glycerol, and 1 mM Mg-GTP in MRB80. The mixture was then immediately placed into 37 ^0^C water bath for 30 min. After polymerization, 10 µM Taxol in MRB80 was added to the polymerized mixture. The mixture was then further incubated in 37 ^0^C water bath for additional 10 min followed by addition of 50 µl 10 µM Taxol in MRB80 (buffer B). The polymerized mix was then centrifuged at 12000 × *g* for 10 min at RT. The supernatant was discarded and the resulting microtubule pellet was resuspended in 50 µl buffer B and was kept in dark for immediate use. GMPCPP stabilized microtubule seeds were assembled in an analogous way by mixing unlabeled, labeled and biotin labeled tubulin 1 mM GMPCPP (Jena Bioscience, # NU405S) followed by incubation on ice for 10 min to allow nucleotide exchange and then polymerizing on the 37 °C water bath for 30 min. 50 µl MRB80 was then added to the polymerization mix and the microtubules were pelleted by centrifugation as above. The resulting microtubule seeds pellet was resuspended in 50 µl MRB80 and stored as above.

#### Synergic binding assay

A microscopy perfusion chamber was prepared by attaching an acid washed coverslip (22 × 22 mm, thickness 1 1/2, Corning) over a pre-cleaned glass slide (75 × 25 mm, thickness 1 mm, Corning) using a double-sided tape (Scotch) resulting in a chamber volume of ∼10 µl. For more details, please see TIRF section in Afreen et al., 2022 [37]. Acid washed coverslips were further cleaned by holding under flame for a couple of seconds prior to use. Solutions were then exchanged into the perfusion chamber by micropipette and filter paper. The chamber was activated by flowing in the following solution in an order in which they appear below, followed by incubation for 5 min and intermittent washing with 3 chamber volumes of buffer B: PLL PEG biotin (0.1 mg ml^-1^, Susos, AG), Streptavidin (0.625 mg ml^-1^, #S4762, Sigma), diluted solution of taxol stabilized microtubules (1 in 100 dilutions form the stock), and κ-Casein (1 mg ml^-1^). All incubations were done inside a wet chamber to prevent evaporation. The chamber was finally washed with imaging buffer (MRB80 supplemented with 10 µM Taxol, 0.6 mg ml^-1^ κ-casein, 4 mM DTT, 50 mM glucose (#G8270, Sigma), 0.2 mg ml^-1^, catalase (#C9322, Sigma), and 0.4 mg ml^-1^glucose oxidase (#G7141, Sigma). Premixed solutions of a GFP labeled protein (either Cdt1, or Ska1, or Ndc80, 1 nM) together with increased concentration untagged proteins (Cdt1, Ska1, or Ndc80, 10 nM – 2 µM) in image buffer were subsequently flown into the chamber and images in were recorded with the following microscopy settings.

Imaging was performed with a Nikon Ti-2 inverted microscope equipped with Photometrics prime 95B sCMOS camera (Teledyne photometrics); 100×, 1.49 NA objective lens. The microscope produced images of pixel size 109 nm, and with excitation laser launcher (Nikon) containing 488, 546, and 640 nm laser lines among other lines. NIS-elements software (Nikon) was used for data acquisition with 100 ms exposure times.

Dynamic experiment was carried out with Nikon Ti-E inverted microscope equipped with Andor iXon3 CCD camera (Cambridge Scientific, Watertown, MA, USA), 1.49× NA, and 100× oil objective. The microscope produced a 512 × 512-pixel images with 0.16 µm per pixel resolution in both *x* and *y* directions. All experiments with dynamic microtubules were carried out at room temperature. The microscope was also equipped with differential interference contrast (DIC) imaging and microtubules were visualized using DIC mode with 100% lamp power and 300 ms exposure time.

#### Single molecule co-localization assay

A microscopy perfusion chamber with immobilized taxol stabilized microtubules was prepared as described above. The chamber was washed with image buffer and microtubule-binding proteins were flown into the chamber. Extremely dilute solutions of GFP-Hec1/Nuf2 dimer (1 nM) and Cy3-Ska1 (1 nM) in image buffer was flown in to chamber together with increasing concentration (0, 1, and 10 nM) Cdt1. For three party colocalization experiments, non-fluorescently labeled microtubule was used together with nanomolar (1 nM) concentrations of GFP-Cdt1, Ska1^Cy3^, and Ndc80^Alexa647^.

#### Experiment with dynamic microtubules

A microscopy perfusion chamber with immobilized GMPCPP stabilized microtubules was prepared as described above. The chamber was washed with image buffer supplemented with 1 mM GTP. A solution of 1 mg/ml tubulin was flown in image buffer (supplemented with 1 mM GTP) together with 1 nM GFP-Cdt1 and 1 nM untagged Ska1.

The chamber was sealed with nail polish and a time series was recorded at room temperature in brightfield-DIC and TIRF mode (488 nm excitation, 100 ms exposure) for every 2 s time interval and for 30 min.

Single molecule diffusion experiment: This method is same as that of the co-localization assays except that the data acquisition was done using 640 nm excitation laser, 40 ms exposure time, 999 EM gain, at 14 MHz read out speed. The data was acquired continually for 2s. Later Kymographs were plotted using kymograph direct software [38] and MSD values were also obtained using the same software. Multiple MSD values were averaged and the averaged MSD vs time plot was analyzed by linear regression to obtain the diffusion coefficient.

### Statistical analysis

Origin software (version 2018) was used for all data plots and obtaining the statistical significance (Mann-Whitney U-test) between data sets.

## Acknowledgments

We thank Dr. Gary Gorbsky (Oklahoma Medical Research Foundation, Oklahoma City) for GFP-SKA1 and GFP-SKA1^ΔMTBD^ expressing stable HeLa cell lines, bacterial expression plasmid encoding GST-tagged SKA3, and for the SKA1 and SKA3 polyclonal antibodies; Dr. A. Arockia Jeyaprakash (University of Edinburgh) for bacterial expression plasmid encoding GFP-SKA2; Dr. P Todd Stukenberg (University of Virginia Charlottesville) for the bacterial expression plasmid encoding His-SKA1/2; Dr. Andrea Musacchio (Max Plank Institute-Dortmund) for the purified human Ndc80 protein complex; Dr. Arshad Desai (University of California San Diego) for the Hec1/Nuf2 dimeric bacterial expression plasmids; Dr. Jeanette Cook (UNC-Chapel Hill) for the valuable intellectual input; Dr. Anita Varma (Northwestern University-Chicago) for technical support and Annie Wang (Northwestern University-Chicago) for help with data analyses.. This work was supported by NIGMS grant R01GM135391 to DV; and NIGMS grant R35GM124889 to RJM. The authors declare no competing financial interests.

---

**Supplementary figure 1:**
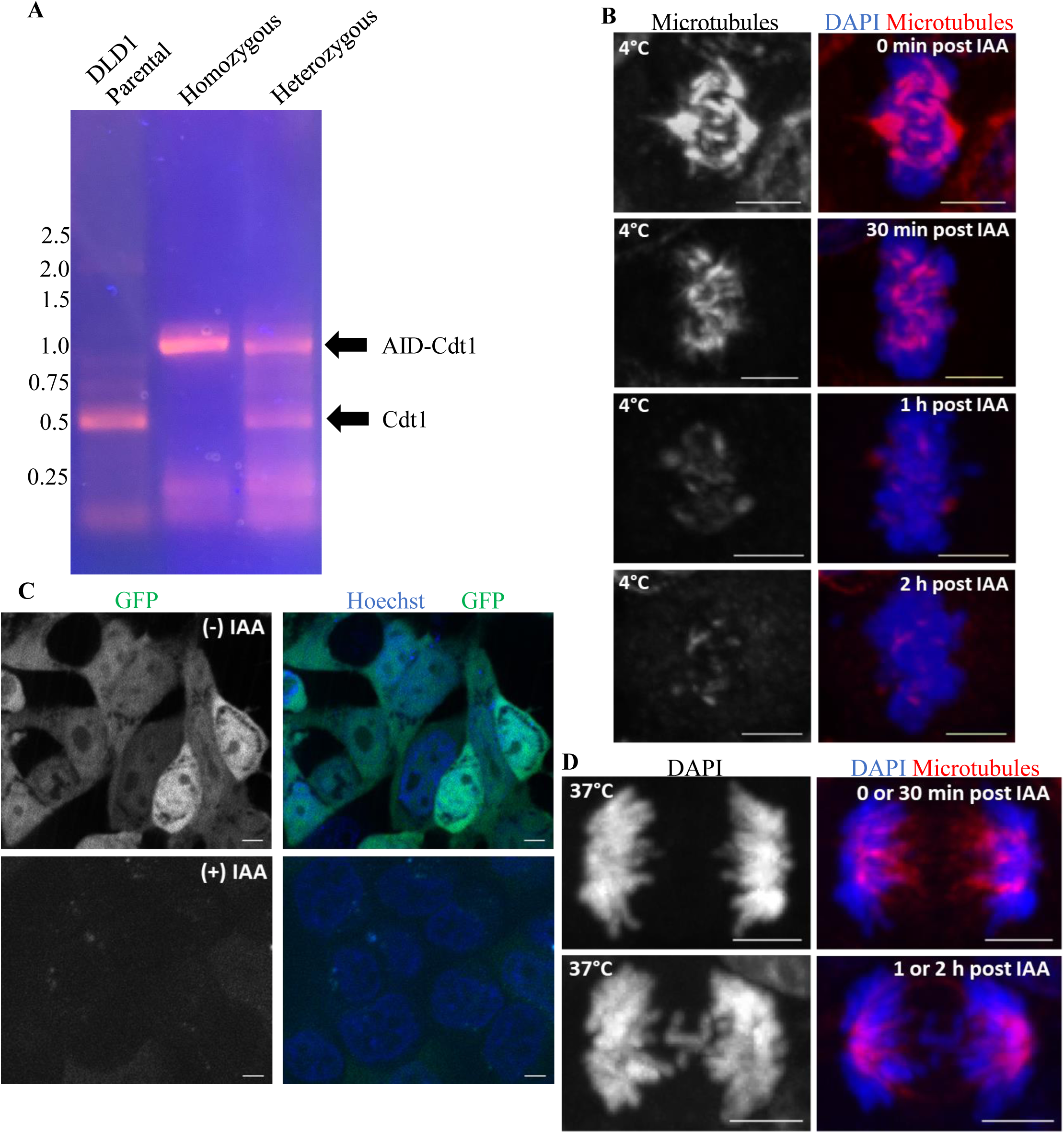
Analyses of the AID-Cdt1 degron system to study mitotic functions of Cdt1. (A) Selection of homozygous AID-Cdt1 colonies by PCR. (B) AID-Cdt1 DLD1 cells growing at 37°C were treated with the DNA stain, Hoechst, along with either control DMSO (top panel) or IAA (bottom panel) for 2 hrs and still pictures of the chromosomes and YFP were acquired. (C) AID-Cdt1 DLD1 cells growing at 37°C were treated with ice-cold buffer for 10 min. after adding IAA for the indicated periods of time. The cells were then immunostaining for antibodies against tubulin and the chromosomes counterstained with DAPI. Scale Bar, 5 µm. (D) AID-Cdt1 DLD1 cells growing at 37°C were fixed after adding IAA for the indicated periods of time. The cells were then immunostaining for antibodies against tubulin and the chromosomes counterstained with DAPI. Scale Bar, 5 µm.

**Supplementary figure 2.**
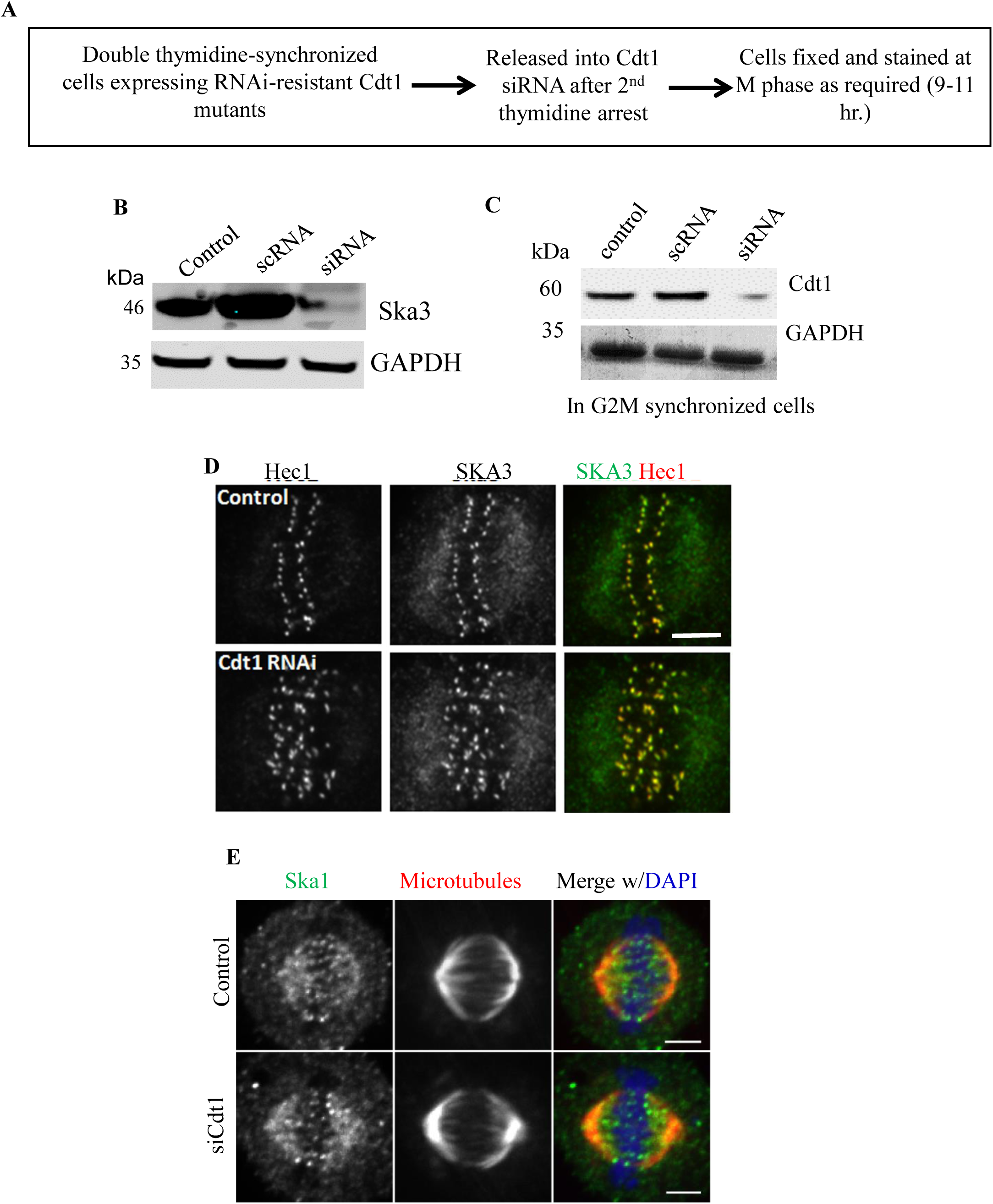
Depletion of Cdt1 from mitotic cells does not interfere with the normal targeting of the Ska1 complex. (A) Schematic representation of the Cdt1 knockdown protocol specifically during mitosis in double thymidine synchronized HeLa cells. (B) Representative Western blots shown with indicated antibodies in each case to assess the level of Ska3 knockdown upon siRNA addition. (C) Same as in B but to assess the level of Cdt1 knockdown upon siRNA addition. (D) HeLa cells treated with either scramble control or siRNA against endogenous Cdt1 were fixed using paraformaldehyde. Representative images of cells immunostained with antibodies against Hec1 (kinetochore marker) in red and SKA3 in green are shown. Scale bar, 5 μm. (E) HeLa cells treated with either scramble control or siRNA against endogenous Cdt1 were fixed using paraformaldehyde. Representative images of cells immunostained with antibodies against SKA3 (in green) and tubulin (in red) are shown; DAPI stained the chromosomes and shown in b/w. Scale bar, 5 μm

**Supplementary figure 3:**
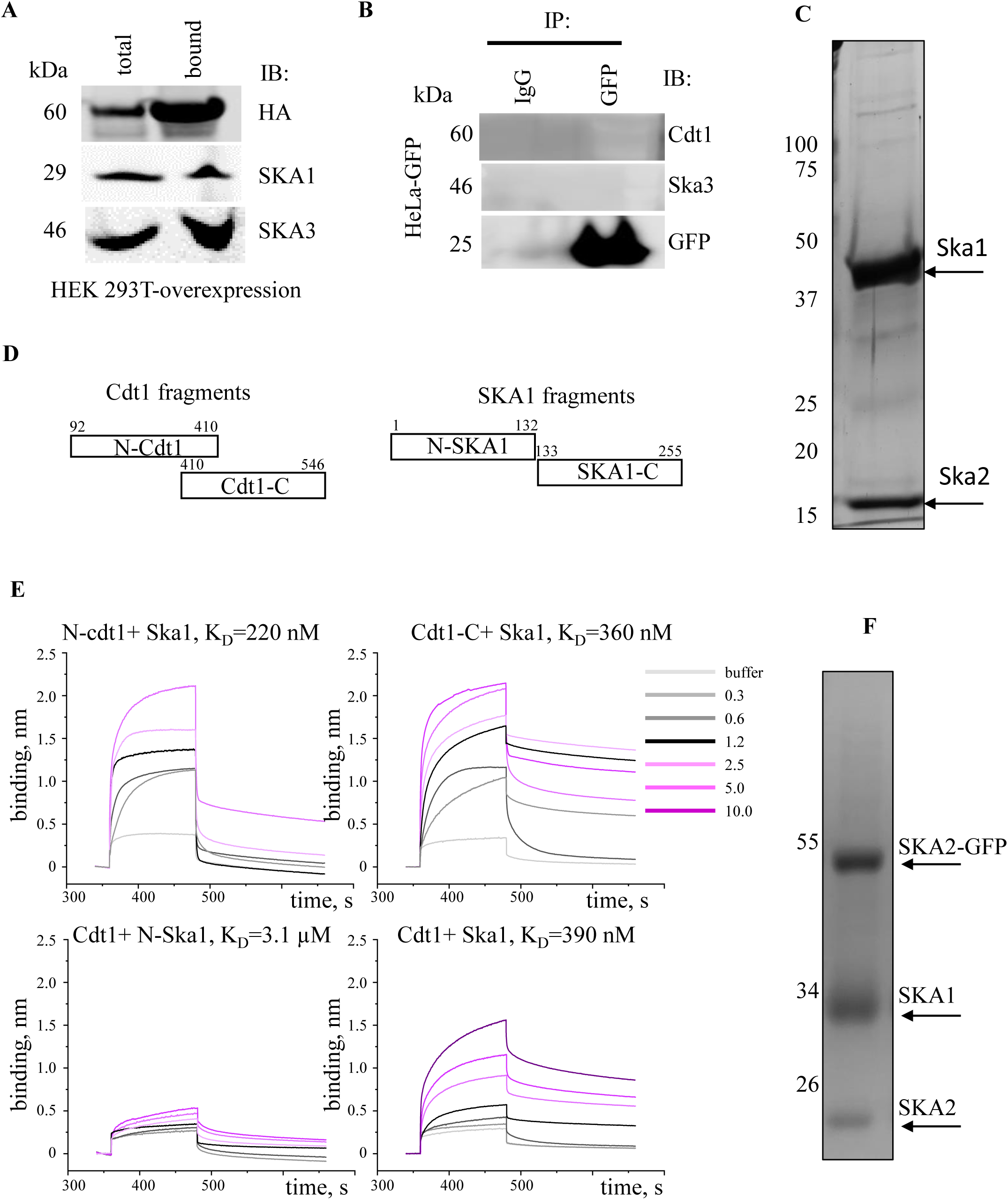
Analyses of binding between Cdt1, Ska1 and their fragments. (A) Pull down of SKA1 and SKA3 by HA/His-tagged Cdt1-WT from thymidine synchronized and nocodazole arrested mitotic HEK-293T cell extracts. The pull down was performed using Ni^+2^-NTA agarose beads followed by immunoblotting with either anti-HA or anti-SKA3 and SKA1 antibodies. 1% of the lysate was loaded as total protein. (B) Thymidine synchronized and nocodazole arrested mitotic HeLa cells stably expressing GFP were immunoprecipitated (IP) and immunoblotted (IB) with the indicated antibodies; IgG was taken as a negative control. 1% of the lysate was loaded as input. (C) SDS-PAGE gel electrophoresis of purified His-SKA1/2 complex (D) Linear protein diagrams of the truncated protein constructs used in the BLI-interaction assay with indicated amino acids position showing truncation sites. (E) BLI sonograms for indicated conditions and their respective K_d_ values. (F) SDS-PAGE gel electrophoresis of purified SKA1/2 +SKA2-GFP complex.

**Supplementary figure 4:**
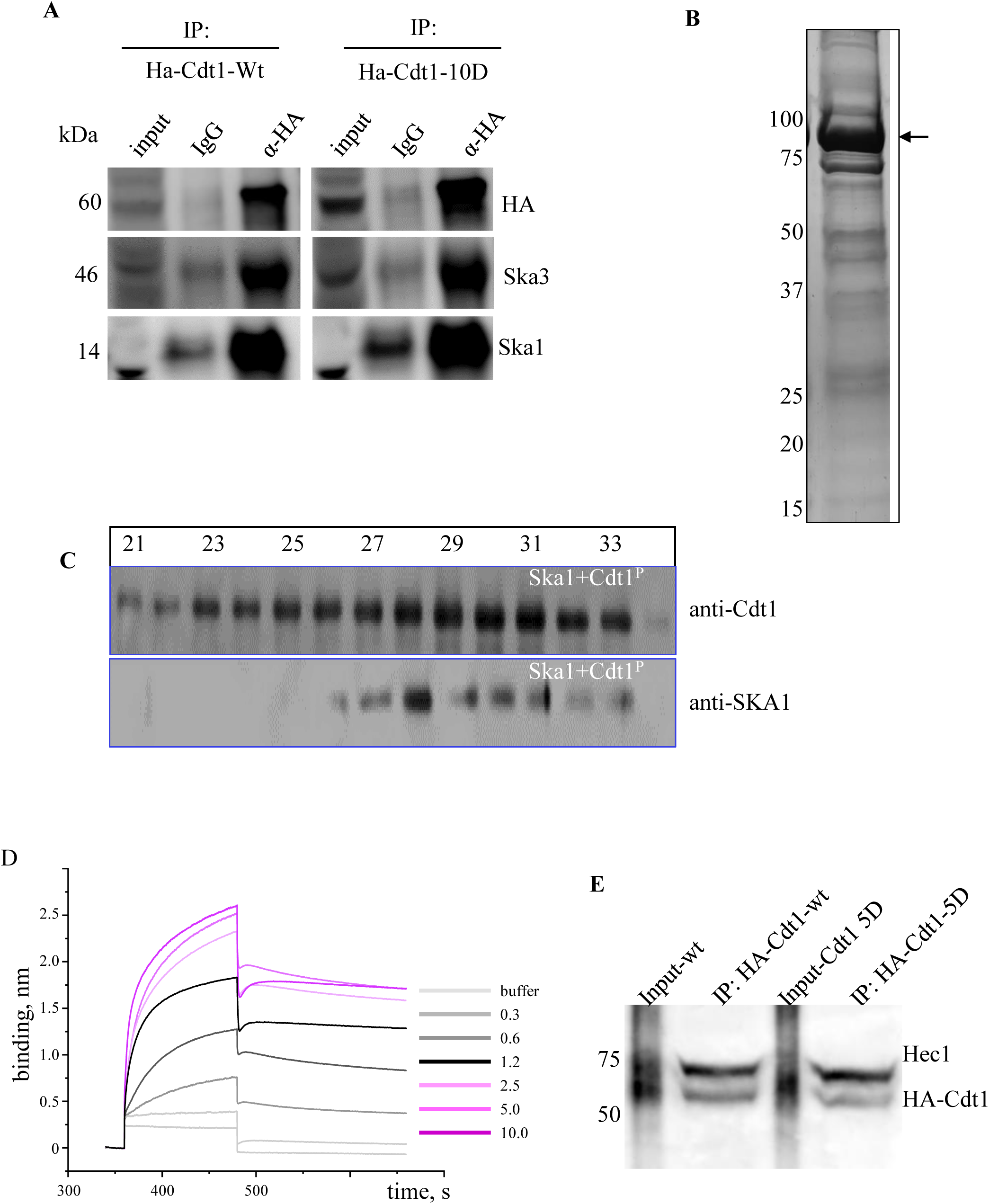
Binding studies of phosphomemeitic Cdt1 and Ska1. (A) Thymidine synchronized and nocodazole arrested mitotic HeLa cells that were stably expressing Cdt1-WT or Cdt1-10D (Aurora B phosphomimetic mutant) were immunoprecipitated (IP) and immunoblotted (IB) with the indicated antibodies; IgG was taken as a negative control. 1% of the lysate was loaded as input. (B) SDS-PAGE gel electrophoresis of purified Cdt1-2E3D protein. (C) Western blot of fractions from co-fractionation experiments for indicated protein complexes probed with anti-Cdt1 (top) and anti SKA1 (bottom). (D) BLI-sensograms showing wavelength shifts (nm) generated by the addition of 0.5 μg/ml Cdt1-2E3D protein with increasing concentrations of SKA1 as indicated in the plot. (E) Nocodazole arrested mitotic HeLa cells that were stably expressing Cdt1-WT or Cdt1-5D (CDK1 phosphomimetic mutant) were immunoprecipitated (IP) with the antibodies indicated on the right.

**Supplementary figure 5:**
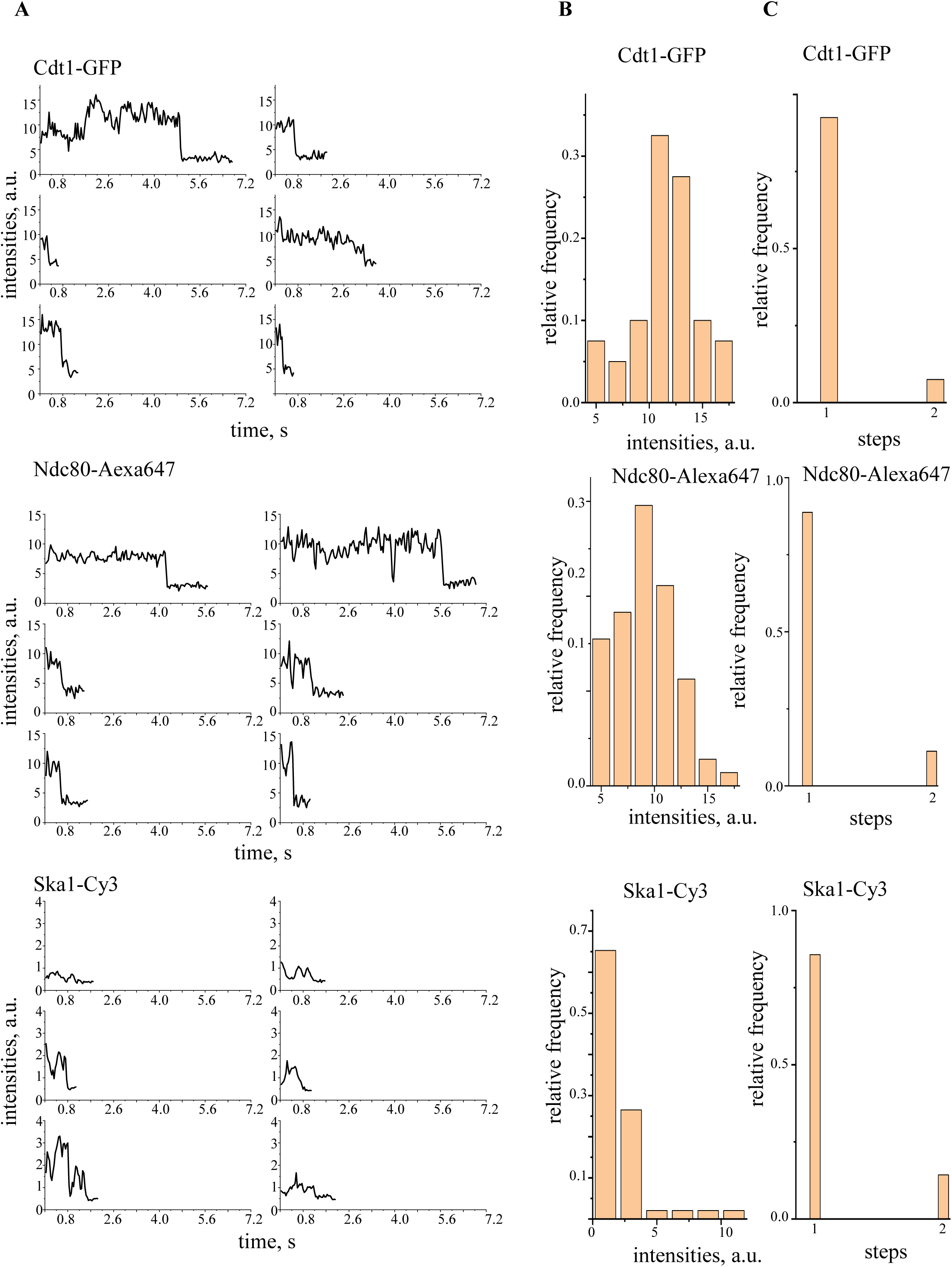
Representative photobleaching steps for GFP-Cdt1, SKA1-Cy3, and Ndc80-Alexa647. A) Representative photobleaching steps for indicated proteins (B) Histogram of starting fluorescence intensity of indicated single molecules and (C) photobleaching steps from n>100 single molecule binding events and N=3 independent trials.

**Supplementary figure 6:**
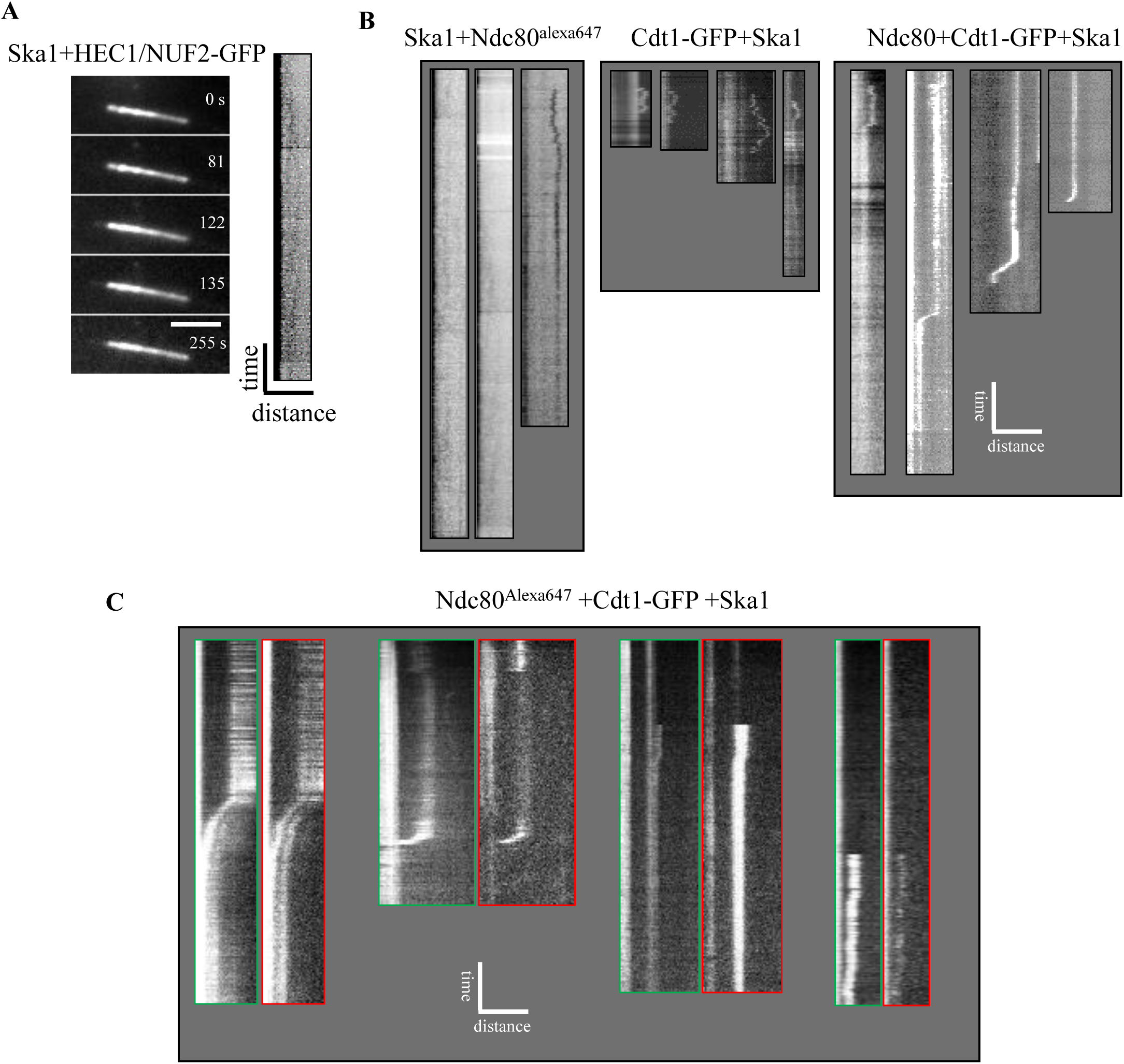
Example kymographs for dynamic microtubules in presence of the indicated combinations of proteins. A) Representative time lapse images recorded with surface immobilized microtubules after addition of a mixture of HEC1/NUF2-GFP, untagged SKA1 (1 nM), soluble tubulin (1 mg/ml), and Mg-GTP (1 mM). Time in s and bar is 5 µm. (B) More example kymographs for dynamic microtubule end tracking by the indicated proteins on top of each panel, bar 1 min., 5 μm. (C) More example kymographs for dynamic microtubule end tracking by GFP-Cdt1, untagged SKA1, and Ndc80^Alexa 647^. Panels showing kymographs for the same end tracking evet is GFP-Cdt1 channel, and in Ndc80^Alexa647^ channel.

## Supplemental Video Legends

**Video 1. Normal mitotic progression in AID-Cdt1 DLD1 cells treated with control DMSO.**

Live images of chromosomes labeled with DNA dye, Hoechst, were captured every 7.5 mins after, for a total period of ∼ 90 min (for this example). The movie was sped up to ∼ 1500 x and played at ∼ 4 frames/s.

**Video 2. Severe delay in mitotic progression in AID-Cdt1 DLD1 cells treated with Auxin, IAA.**

Live images of chromosomes labeled with DNA dye, Hoechst, were captured every 7.5 mins after, for a total period of ∼ 350 min (for this example). The movie was sped up to ∼ 1750 x and played at ∼ 4 frames/s.

**Video 3. Chromosome mis-segregation in AID-Cdt1 DLD1 cells treated with Auxin, IAA.**

Live images of chromosomes labeled with DNA dye, Hoechst, were captured every 7.5 mins after, for a total period of 135 min (for this example). The movie was sped up to ∼ 1600 x and played at ∼ 4 frames/s.

**Video 4. GFP-Cdt1 diffusively tracks the end of dynamic microtubule in presence of Ska1.**

Video shows GMPCPP stabilized microtubules decorated with GFP-Cdt1 visualized with 488 nm excitation laser in TIRF. A complex of GFP-Cdt1 and Ska1 (not imaged) lands near the growing end of microtubule1 at about 1 s and starts diffusively tracking the dynamic end till 12 s, then it detaches and diffuses into the solution out of field of view. Movie is played at 30 times faster rate than real. Scale bar 3 µm.

**Video 5. Ndc80-Alexa 647 diffusively tracks the end of dynamic microtubule in presence of Ska1.**

Video shows GMPCPP stabilized microtubules decorated with Ndc80^alexa 647^ visualized with 640 nm excitation laser in TIRF. A very weak intensity of Ndc80^alexa 647^ seen near the dynamic end for the entire duration of the movie. Movie is played at 60 times faster rate than real. Scale bar 3 µm.

**Video 6. GFP-Cdt1 robustly tracks the end of dynamic microtubule in presence of Ndc80 and Ska1.**

Video shows GMPCPP stabilized microtubules decorated with GFP-Cdt1 visualized with 488 nm excitation laser in TIRF. A complex of GFP-Cdt1, Ska1 and Ndc80 (not imaged) already bound to the grown end of a microtubule at the beginning of the movie and starts to robustly track the depolymerizing phase of the dynamic end at 1 s, then it remains bound on the microtubule lattice for the entire duration of the movie. Movie is played at 30 times faster rate than real. Scale bar 3 µm.

**Video 7. GFP-Cdt1/ Ndc80^Alexa647^ robustly tracks the end of dynamic microtubule in presence of Ska1.**

Video shows GMPCPP stabilized microtubules decorated with GFP-Cdt1 and Ndc80^alexa 647^ visualized with 488/640 nm excitation lasers in TIRF. A complex of GFP-Cdt1, Ndc80^alexa 647^, and Ska1 (not imaged) is seen to track the dynamic end of microtubules for couple of cycles of polymerization and depolymerization. Movie is played at 30 times faster rate than real. Scale bar 3 µm.

## Notes

### Competing Interest Statement

The authors have declared no competing interest.

## References

1. Pozo, P.N., and Cook, J.G. (2016). Regulation and Function of Cdt1; A Key Factor in Cell Proliferation and Genome Stability. Genes (Basel) 8.

2. Cheeseman, I.M., and Desai, A. (2008). Molecular architecture of the kinetochore-microtubule interface. Nat Rev Mol Cell Biol 9, 33–46.

3. Varma, D., Chandrasekaran, S., Sundin, L.J., Reidy, K.T., Wan, X., Chasse, D.A., Nevis, K.R., DeLuca, J.G., Salmon, E.D., and Cook, J.G. (2012). Recruitment of the human Cdt1 replication licensing protein by the loop domain of Hec1 is required for stable kinetochore-microtubule attachment. Nat Cell Biol 14, 593–603.

4. Guimaraes, G.J., Dong, Y., McEwen, B.F., and Deluca, J.G. (2008). Kinetochore-microtubule attachment relies on the disordered N-terminal tail domain of Hec1. Curr Biol 18, 1778–1784.

5. Cheeseman, I.M., Chappie, J.S., Wilson-Kubalek, E.M., and Desai, A. (2006). The conserved KMN network constitutes the core microtubule-binding site of the kinetochore. Cell 127, 983–997.

6. Shivangi Agarwal, K.P.S., Yizhuo Zhou, Aussie Suzuki, Richard J. McKenney, Dileep Varma (2018). Human replication licensing factor Cdt1 binds spindle microtubules and is a substrate for Aurora B kinase Journal of Cell Biology

7. Hanisch, A., Sillje, H.H., and Nigg, E.A. (2006). Timely anaphase onset requires a novel spindle and kinetochore complex comprising Ska1 and Ska2. EMBO J 25, 5504–5515.

8. Welburn, J.P., Grishchuk, E.L., Backer, C.B., Wilson-Kubalek, E.M., Yates, J.R., 3rd, and Cheeseman, I.M. (2009). The human kinetochore Ska1 complex facilitates microtubule depolymerization-coupled motility. Dev Cell 16, 374–385.

9. Schmidt, J.C., Arthanari, H., Boeszoermenyi, A., Dashkevich, N.M., Wilson-Kubalek, E.M., Monnier, N., Markus, M., Oberer, M., Milligan, R.A., Bathe, M., et al. (2012). The kinetochore-bound Ska1 complex tracks depolymerizing microtubules and binds to curved protofilaments. Dev Cell 23, 968–980.

10. Jeyaprakash, A.A., Santamaria, A., Jayachandran, U., Chan, Y.W., Benda, C., Nigg, E.A., and Conti, E. (2012). Structural and functional organization of the Ska complex, a key component of the kinetochore-microtubule interface. Mol Cell 46, 274–286.

11. Zhang, G., Kelstrup, C.D., Hu, X.W., Kaas Hansen, M.J., Singleton, M.R., Olsen, J.V., and Nilsson, J. (2012). The Ndc80 internal loop is required for recruitment of the Ska complex to establish end-on microtubule attachment to kinetochores. J Cell Sci 125, 3243–3253.

12. Powers, A.F., Franck, A.D., Gestaut, D.R., Cooper, J., Gracyzk, B., Wei, R.R., Wordeman, L., Davis, T.N., and Asbury, C.L. (2009). The Ndc80 kinetochore complex forms load-bearing attachments to dynamic microtubule tips via biased diffusion. Cell 136, 865–875.

13. Chan, Y.W., Jeyaprakash, A.A., Nigg, E.A., and Santamaria, A. (2012). Aurora B controls kinetochore-microtubule attachments by inhibiting Ska complex-KMN network interaction. J Cell Biol 196, 563–571.

14. Raaijmakers, J.A., Tanenbaum, M.E., Maia, A.F., and Medema, R.H. (2009). RAMA1 is a novel kinetochore protein involved in kinetochore-microtubule attachment. J Cell Sci 122, 2436–2445.

15. Chandrasekaran, S., Tan, T.X., Hall, J.R., and Cook, J.G. (2011). Stress-stimulated mitogen-activated protein kinases control the stability and activity of the Cdt1 DNA replication licensing factor. Mol Cell Biol 31, 4405–4416.

16. Chakraborty, M., Tarasovetc, E.V., and Grishchuk, E.L. (2018). Chapter 13 - In vitro reconstitution of lateral to end-on conversion of kinetochore–microtubule attachments. In Methods in Cell Biology, Volume 144, H. Maiato and M. Schuh, eds. (Academic Press), pp. 307–327.

17. Huis In ’t Veld, P.J., Volkov, V.A., Stender, I.D., Musacchio, A., and Dogterom, M. (2019). Molecular determinants of the Ska-Ndc80 interaction and their influence on microtubule tracking and force-coupling. Elife 8.

18. Rahi, A., Chakraborty, M., Vosberg, K., and Varma, D. (2020). Kinetochore–microtubule coupling mechanisms mediated by the Ska1 complex and Cdt1. Essays in Biochemistry 64, 337–347.

19. Monda, J.K., and Cheeseman, I.M. (2018). The kinetochore-microtubule interface at a glance. J Cell Sci 131.

20. Gaitanos, T.N., Santamaria, A., Jeyaprakash, A.A., Wang, B., Conti, E., and Nigg, E.A. (2009). Stable kinetochore-microtubule interactions depend on the Ska complex and its new component Ska3/C13Orf3. EMBO J 28, 1442–1452.

21. Daum, J.R., Wren, J.D., Daniel, J.J., Sivakumar, S., McAvoy, J.N., Potapova, T.A., and Gorbsky, G.J. (2009). Ska3 is required for spindle checkpoint silencing and the maintenance of chromosome cohesion in mitosis. Curr Biol 19, 1467–1472.

22. Varma, D., and Salmon, E.D. (2012). The KMN protein network--chief conductors of the kinetochore orchestra. J Cell Sci 125, 5927–5936.

23. Abad, M.A., Medina, B., Santamaria, A., Zou, J., Plasberg-Hill, C., Madhumalar, A., Jayachandran, U., Redli, P.M., Rappsilber, J., Nigg, E.A., et al. (2014). Structural basis for microtubule recognition by the human kinetochore Ska complex. Nature Communications 5, 2964.

24. Abad, M.A., Zou, J., Medina-Pritchard, B., Nigg, E.A., Rappsilber, J., Santamaria, A., and Jeyaprakash, A.A. (2016). Ska3 Ensures Timely Mitotic Progression by Interacting Directly With Microtubules and Ska1 Microtubule Binding Domain. Sci Rep 6, 34042.

25. Hill, T.L. (1985). Theoretical problems related to the attachment of microtubules to kinetochores. Proc Natl Acad Sci U S A 82, 4404–4408.

26. Koshland, D.E., Mitchison, T.J., and Kirschner, M.W. (1988). Polewards chromosome movement driven by microtubule depolymerization in vitro. Nature 331, 499–504.

27. Campbell, S., Amin, M.A., Varma, D., and Bidone, T.C. (2019). Computational model demonstrates that Ndc80-associated proteins strengthen kinetochore-microtubule attachments in metaphase. Cytoskeleton 76, 549–561.

28. Saurin, A.T. (2018). Kinase and Phosphatase Cross-Talk at the Kinetochore. Frontiers in Cell and Developmental Biology 6.

29. Moura, M., and Conde, C. (2019). Phosphatases in Mitosis: Roles and Regulation. Biomolecules 9, 55.

30. Gavet, O., and Pines, J. (2010). Progressive activation of CyclinB1-Cdk1 coordinates entry to mitosis. Dev Cell 18, 533–543.

31. Zhou, Y., Pozo, P.N., Oh, S., Stone, H.M., and Cook, J.G. (2020). Distinct and sequential re-replication barriers ensure precise genome duplication. PLoS Genet 16, e1008988.

32. Chakraborty, M., Toleikis, A., Siddiqui, N., Cross, R.A., and Straube, A. (2020). Activation of cytoplasmic dynein through microtubule crossbridging. bioRxiv, 2020.2004.2013.038950.

33. Li, S., Prasanna, X., Salo, V.T., Vattulainen, I., and Ikonen, E. (2019). An efficient auxin-inducible degron system with low basal degradation in human cells. Nat Methods 16, 866–869.

34. Cong, L., Ran, F.A., Cox, D., Lin, S., Barretto, R., Habib, N., Hsu, P.D., Wu, X., Jiang, W., Marraffini, L.A., et al. (2013). Multiplex genome engineering using CRISPR/Cas systems. Science 339, 819–823.

35. Zasadzińska, E., Huang, J., Bailey, A.O., Guo, L.Y., Lee, N.S., Srivastava, S., Wong, K.A., French, B.T., Black, B.E., and Foltz, D.R. (2018). Inheritance of CENP-A Nucleosomes during DNA Replication Requires HJURP. Dev Cell 47, 348–362.e347.

36. Muller-Esparza, H., Osorio-Valeriano, M., Steube, N., Thanbichler, M., and Randau, L. (2020). Bio-Layer Interferometry Analysis of the Target Binding Activity of CRISPR-Cas Effector Complexes. Front Mol Biosci 7, 98.

37. Afreen, S., Rahi, A., Landeros, A.G., Chakraborty, M., McKenney, R.J., and Varma, D. (2022). In Vitro and In Vivo Approaches to Study Kinetochore-Microtubule Attachments During Mitosis. Methods Mol Biol 2415, 123–138.

38. Mangeol, P., Prevo, B., and Peterman, E.J.G. (2016). KymographClear and KymographDirect: two tools for the automated quantitative analysis of molecular and cellular dynamics using kymographs. Molecular Biology of the Cell 27, 1948–1957.

